# Molecular basis for potent B cell responses to antigen displayed on particles of viral size

**DOI:** 10.1101/2023.02.15.528761

**Authors:** Jeremy F. Brooks, Julianne Riggs, James L. Mueller, Raisa Mathenge, Wei-Yun Wholey, Sekou-Tidiane Yoda, Vivasvan S. Vykunta, Wei Cheng, Julie Zikherman

**Author notes:** Both authors contributed equally to this work (Jeremy F. Brooks, Julianne Riggs). Corresponding authors: Dr Julie Zikherman 513 Parnassus Avenue Room HSW1201E, Box 0795 San Francisco, CA, 94143-0795 email address Dr Wei Cheng University of Michigan 428 Church Street Ann Arbor, MI, 48109-1065.

## Abstract

Although it has long been appreciated that multivalent antigens – and particularly viral epitope display – produce extremely rapid, robust, and T-independent humoral immune responses, the biochemical basis for such potency has been incompletely understood. Here we take advantage of a set of neutral liposomes of viral size that are engineered to display affinity mutants of the model antigen (Ag) hen egg lysozyme at precisely varied density. We show that particulate Ag display by liposomes induces highly potent B cell responses that are dose-and density-dependent but affinity-independent. Titrating dose of particulate, but not soluble, Ag reveals bimodal Erk phosphorylation and cytosolic calcium increases. Particulate Ag induces signal amplification downstream of the B cell receptor (BCR) by selectively evading LYN-dependent inhibitory pathways, but *in vitro* potency is independent of CD19. Importantly, Ag display on viral-sized particles signals independently of MYD88 and IRAK1/4, but activates NF-*κ*B robustly in a manner that mimics T cell help. Together, such biased signaling by particulate Ag promotes MYC expression and reduces the threshold required for B cell proliferation relative to soluble Ag. These findings uncover a molecular basis for highly sensitive B cell response to viral Ag display and remarkable potency of virus-like particle vaccines that is not merely accounted for by avidity and BCR cross-linking, and is independent of the contribution of B cell nucleic acid-sensing machinery.

## Introduction

Multivalent antigens, including viral structures, produce robust B cell responses that can engage – but do not require – T cell help, and design of virus-like particle (VLP) vaccines is predicated in part upon this observation^1, 2^. Particulate HIV immunogens with high valency can recruit precursor B cells expressing rare and low-affinity germline-encoded B cell receptors (BCRs) into both the extra-follicular and germinal center responses^3–5^. Indeed, highly organized epitope display on the surface of viral particles has long been proposed to function as an immunogenic signal of ‘foreignness,’ and can trigger robust T-independent antibody responses that may not require TLR engagement^1, 2, 6–11^.

Although the structural basis for BCR triggering by various forms of antigen (Ag) continues to be debated^12^, potent cellular responses to multivalent epitopes are assumed to reflect cross-linking of Ag-receptors. Dintzis et al. reported that a minimum number of haptens (>12) spaced appropriately along a linear backbone (*≅* 10 nm) was essential to trigger a T-independent humoral immune response *in vivo* and proposed the “immunon” model, postulating a requirement for clustering of a minimum number of hapten-binding receptors^8^. More recently, “molecular origami” has defined an optimal spacing and valency of high affinity epitopes to trigger BCR-induced calcium entry^13^. However, the mechanistic basis for potent (and T-independent) B cell responses to density of epitopes displayed on particles of viral size is not fully understood. Specifically, whether such potency is purely a function of avidity and receptor cross-linking, or also reflects a qualitatively unique mode of B cell triggering by this biologically significant class of Ag is unknown.

By contrast, the molecular basis for B cell responses to membrane-associated Ag presented on the surface of cells has been extensively explored^14–18^. Tethering of non-stimulatory monovalent haptens to a membrane is sufficient to trigger BCR signaling^19^. Membrane-associated Ag also dramatically reduces the threshold for B cell activation relative to soluble Ag, and impacts affinity discrimination^20–24^. Membrane Ags have been postulated to achieve such potency by reorganizing membrane structure upon contact with B cells to generate initially BCR microclusters, and subsequently an immune-synapse-like (IS) contact interface via actin remodeling^14, 15, 20, 23^. This is associated with recruitment of CD19 co-receptor into the IS and exclusion of inhibitory co-receptors such as CD22 and Fc*γ*RIIb along with associated phosphatase (PTPase) SHP1^20, 25, 26^. Membrane-associated Ags require CD19 for downstream signaling *in vitro* (as does B cell signaling triggered by actin depolymerization), while soluble Ag does not^23, 27–29^. Whether exclusion from the IS of inhibitory machinery is also essential for potent responses to membrane Ag is less clear_20,25,26,30_.

However, Ag captured and presented on the membrane of macrophages and dendritic cells, and particles of viral size differ from one another not only in size of interface presented to responding B cells but also in fundamental biophysical properties such as actin dynamics, pulling force, and BCR internalization. B cells encounter and respond to membrane-associated Ag in a variety of physiologic contexts, such as when Ag is captured and presented on the surface of follicular dendritic cells during affinity maturation. Direct encounter between B cells and viruses – as well as multivalent VLP vaccines – occurs *in vivo* at early time points and the nature of this interaction can define the trajectory of the subsequent immune response^1, 31–37^.

To understand how B cells signal in response to Ag display on viruses, here we take advantage of a novel set of viral-sized liposomes that are engineered to display affinity mutants of the model Ag hen egg lysozyme (HEL) but lacking nucleic acid cargo^38^. These liposomes display a programmable range of HEL epitope density on their surface that mimics the spectrum observed on naturally occurring viruses^39^. We show that Ag display on these synthetic particles of viral size induces potent and remarkably prolonged B cell responses that are density-dependent but affinity-independent, and do not rely on MYD88/IRAK1/4 signaling. These responses are characterized by bimodal Erk phosphorylation and cytosolic calcium entry at the single cell level suggesting digital behavior, and are sufficient to activate anergic B cells. Signaling in response to these particles requires SYK, BTK, and PI3K enzyme activity, implicating the canonical BCR signaling pathway. We show that such particulate Ags trigger robust signal amplification downstream of the BCR by evading LYN-dependent inhibitory pathways, but *in vitro* potency is independent of CD19 – in contrast to membrane Ag presented on APCs. Particulate Ag also trigger uniquely robust NF-*κ*B activation, MYC expression, cell growth, survival, and reduce the threshold for B cell proliferation in a manner that mimics T cell help. This reveals a new mechanism by which B cells recognize virus-like Ag display as stand-alone danger signal – independent of nucleic acid cargo – that does not rely exclusively on avidity and BCR cross-linking. This in turn helps explain how extremely rapid, protective, and T-independent B cell responses to low doses of virus or VLP are mounted.

## Results

### Antigen display on particles of viral size induce potent B cell responses that are density-dependent but affinity-independent

We constructed a library of neutral, unilamellar liposomes of viral size (approx. 120nm diameter) conjugated via maleimide chemistry to a very well-characterized model Ag (hen egg lysozyme; HEL), as previously described by us (**Figures 1A, Supplementary Figures 1A, B)**^38^. These liposomes display HEL protein in a specific orientation and at a programmable density in order to mimic the range of epitope display on bona fide viruses^39^. Importantly, to isolate the impact of virus-like Ag display on B cell responses, the synthetic virus-like structures (SVLS) generated for this study harbor phosphate-buffered saline on the interior and lack any packaged nucleic acid content such that conjugated surface proteins are the only immunogenic component.

**Figure 1.**
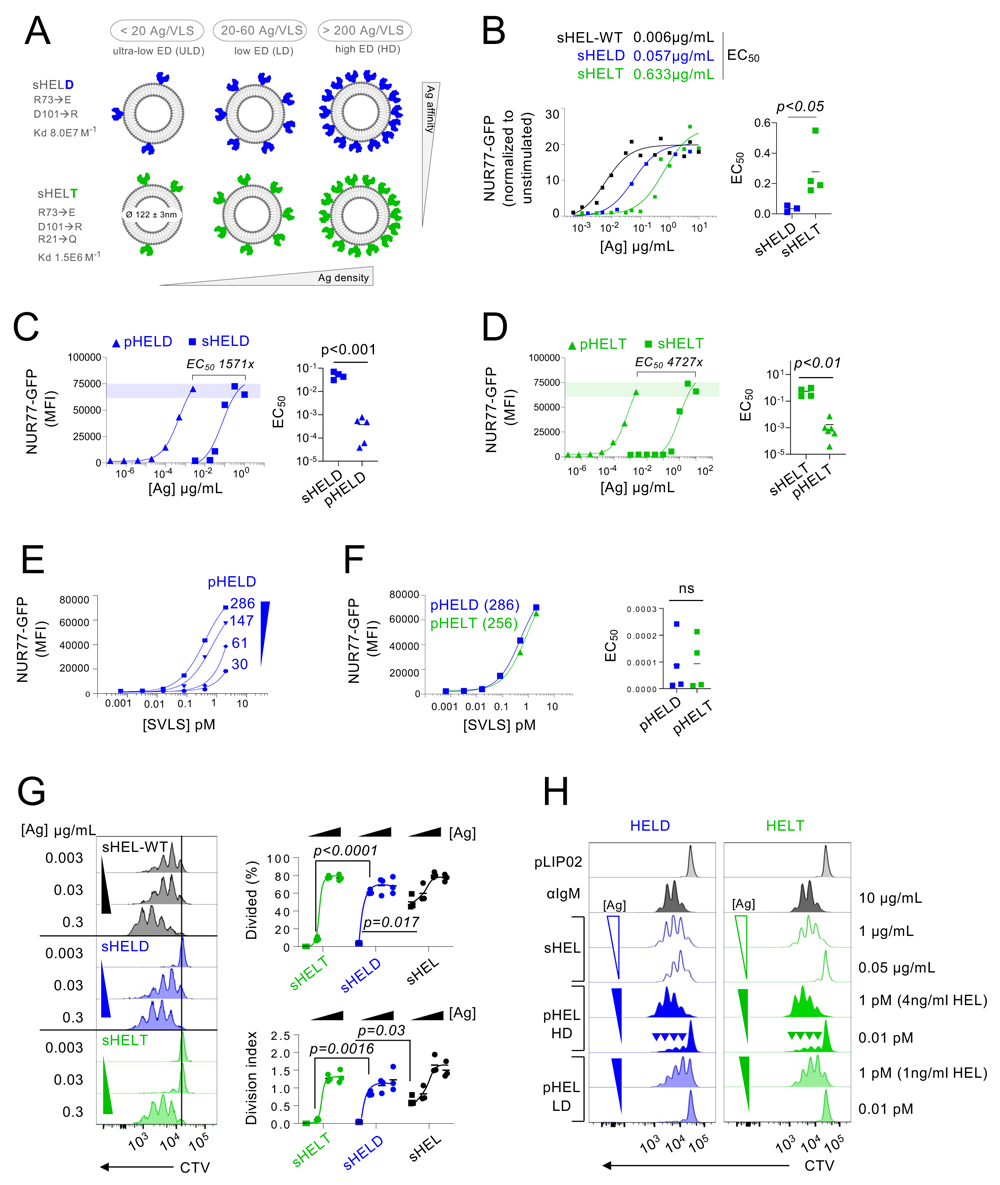
Synthetic virus-like structures (SVLS) induce highly potent B cell responses that depend on epitope density but not affinity. **A.** Library of synthetic viral-like structures (SVLS) assembled by conjugated lipid nanoparticles of viral size (120nm) via maleimide chemistry to engineered cysteines in hen egg lysozyme (HEL) affinity mutants at varying epitope density, ED (see also S1B for quantification). **B.** MD4 lymphocytes expressing NUR77-eGFP BAC Tg reporter were stimulated with native soluble HEL (sHEL-WT) or the affinity mutants sHELD and sHELT at varying doses for 24 hrs. GFP expression was analysed by flow cytometry in B220+ B cells and MFI is plotted. Curves fit a three-parameter non-linear regression against the mean of 3 biological replicates per stimulus and EC50 values correspond to these curves (left). EC50 values were also interpolated for individual biological replicates and compared by parametric t-test (right). **C.** Pooled splenocytes and lymphocytes from NUR77-GFP reporter MD4 mice were stimulated with either soluble HELD (sHELD) or SVLS carrying 286 HELD epitopes (pHELD) and analysed 24 hrs later as in B. Data from a representative experiment (left panel) were fitted with a three-parameter non-linear regression. Shaded bar corresponds to equipotent doses of sHELD (1 μg/ml) and pHELD (1pM ∼ 4ng/ml HEL). EC50 values were interpolated from 4-5 biological replicates and compared by parametric t-test (right panel). **D.** As in C, but MD4 reporter B cells were stimulated with soluble HELT (sHELT) and SVLS carrying 256 HELT epitopes (pHELT). **E.** As in C, but MD4 reporter B cells were stimulated with titrated densities of HELD per particle. **F.** pHELT and pHELD curves from C, D are overlaid on the same plot (left). EC50 values [μg/ml] HEL protein were interpolated from 4 biological replicates and compared by parametric t-test (right panel). Data in C-F are representative of 7 independent experiments. **G.** MD4 lymphocytes were loaded with CTV vital dye and cultured with soluble HEL stimuli (sHELT, sHELD, sHEL-WT; 3.16 μg/ml and 10-fold dilutions x 4) together with 20 ng/ml BAFF. Representative histograms depict CTV dilution in live B220+ cells after 72 hours (Left). Graphs depict % divided and division index calculated with Flowjo software for 3 biological replicates (right). Groups were compared by parametric two-way ANOVA with Tukey’s. **H.** MD4 lymph node cells were loaded with CTV vital dye and cultured with either sHELD/T 1 μg/ml, 0.05 μg/ml or pHELT/D with high or low ED (pHELD-HD = 286, pHELT-HD = 256, pHELD-LD = 61, pHELT-LD = 64) at 1pM or 0.1pM. Representative histograms depict CTV dilution in live B220+ cells after 72 hours. Inset triangles denote high and low dilution of stimulus. Inset arrowheads mark a small fraction of highly proliferative B cells. Histograms are representative of 7 independent experiments.

We further took advantage of previously characterized amino acid substitutions in the HEL protein that reduce its affinity for the Hy10 BCR relative to native (“WT”) HEL ^40^. HELD refers to a HEL mutant carrying double mutations R73E, D101R (Ka to the Hy10 BCR = 8.0×10^7^ M^−1^) and HELT refers to a HEL mutant carrying triple mutations R21Q, R73E, D101R (Ka = 1.5×10^6^ M^−1^) **(Supplementary Figures 1A)**^40^. Importantly, the micromolar affinity of the mutant HELT protein for the Hy10 BCR is comparable to typical germline-encoded naïve B cell affinities for polyclonal Ags^3^. This feature enables us to probe physiologically relevant interactions and to extrapolate our findings to a naturally occurring naïve B cell repertoire. We confirmed that the soluble HEL mutant proteins we generated triggered responses in an affinity-and concentration-dependent manner in naïve B cells from so-called “MD4” mice that express the Hy10 IgM/IgD BCR transgene and co-express a reporter of BCR signaling, NUR77/*Nr4a1*-eGFP BAC Tg^41, 42^ **(Figure 1B),** consistent with previous reports^38, 40, 43^.

By combining engineered liposomes with recombinant HEL proteins, we can vary both density and affinity of conjugated surface epitopes (**Figures 1A, Supplementary Figures 1A, B).** We decorated liposomes with either the moderate affinity mutant (HELD) or the low affinity mutant (HELT) at matched densities, between 3-500 epitopes per liposomal particle (approximately 60 to 10,000 molecules per µm^2^); these HEL-engineered SVLS are referred to herein as particulate HEL or pHEL – by contrast to soluble HEL or sHEL **(Figure 1A, Supplementary Figure 1B)**.

The concentration of HELD and HELT proteins required to upregulate NUR77-GFP in MD4 reporter B cells expressing the Hy10 BCR was dramatically reduced (by over 3 orders of magnitude) when tethered to SVLS at a density of approximately 250-290 epitopes per liposome **(Figure 1C, D)**, termed epitope density or ED herein. For example, the EC50 value for sHELT in this assay was on the order of 500 ng/ml, consistent with prior reports^38, 40, 43^, but nearly 5000-fold lower concentrations of HELT protein when tethered to SVLS (pHELT) could elicit detectable eGFP upregulation in Ag-specific reporter B cells after 24 hours of *in vitro* culture. Interestingly, a comparable 1000-fold boost in potency was previously described for intermediate affinity HEL (Ka similar to HELD mutant in our study) tethered to a cell membrane^20^. Non-transgenic (wild-type) NUR77-GFP reporter B cells admixed with MD4 reporter B cells in culture failed to upregulate GFP in response to pHEL confirming that Ag-specific BCR was required for the response **(Supplementary Figure 1C)**. Similarly, the response was not simply due to the physical properties of the particles themselves, because “naked” liposomes (pLIP02) without conjugated HEL protein also failed to induce NUR77-GFP in MD4 B cells **(Supplementary Figure 1C)**. Impressively, a single batch of pHEL SVLS stored at 4 degrees was stable for more than 400 days following its synthesis without loss of potency **(Supplementary Figure 1D)**.

To study effects of ED among engineered liposomes, pHEL library constructs were compared according to liposome concentration across a broad titration in order to control for liposome number. B cells are highly sensitive to ED of pHELD and upregulated NUR77-GFP in a density-dependent manner **(Figure 1E)**. Interestingly, B cell activation did not scale continuously with ED for low affinity pHELT **(Supplementary Figure 1E)**, indicating that there may be an optimum ED for display that is not simply the highest. Moreover, even particles with as few as 3 conjugated HELT molecules could trigger NUR77-GFP upregulation at particle concentration < 1pM **(Supplementary Figure 1F)**, suggesting that tethering of Ag to SVLS even in the absence of extensive BCR cross-linking confers many orders of magnitude higher potency than soluble Ag (**Figure 1B)**. Similar results were obtained with an orthogonal readout of B cell activation, surface CD69 upregulation, and also segregated by high and low ED **(Supplementary Figure 1G)**. B cell sensitivity to ED was also observed *in vivo* (doi: https://doi.org/10.1101/2023.02.20.529089).

Prior work has demonstrated that conjugating similar HEL Ags to large cell surface membranes can override affinity discrimination^21, 22^, although B cells still exhibit affinity-sensing behavior in response to Ag presented on cells^23, 24^. Here we show that conjugating HEL affinity mutants to SVLS also bridges affinity gaps; SVLS bearing either HELD or HELT Ag at matched ED were capable of inducing equivalent levels of NUR77-GFP expression despite a reproducible gap in affinity and potency measured for these proteins in soluble form **(Figure 1B, F, Supplementary Figure 1F)**, indicating that B cells could be activated even by low affinity Ag displayed on small particles of viral size.

Collectively our data demonstrate that Ag display on SVLS are highly potent for inducing B cell responses in a density-dependent but affinity-independent manner. Importantly, these 24 hr assays enabled us to select “equipotent” doses of pHEL and sHEL for further study of both upstream and downstream B cell responses. Specifically, 1pM pHELD/pHELT liposomes conjugated with high ED (250-290 epitopes / SVLS) corresponds to 250-290pM (*≅* 4ng/ml) of HEL protein. Similarly, 1pM of lower ED pHEL liposomes (61-64 epitopes / SVLS) corresponds to 61-64pM (*≅* 1ng/ml) HEL protein (**Supplementary Figure 1B**). Throughout this manuscript, 1pM pHEL doses were compared to much higher, maximally potent sHEL concentrations of 1µg/ml (*≅* 64nM HEL protein) except where dose is explicitly titrated (0.1-10pM range). Indeed, high doses of soluble stimuli – especially low affinity mutants of HEL – are necessary to trigger detectable B cell signaling and activation. These ‘equipotent’ doses used for subsequent assays therefore corresponded to 250-1000x differences in HEL concentration between pHEL and sHEL comparators depending on ED (**Figures 1C, D**).

### Low doses of pHEL drive robust B cell proliferation in an affinity-independent manner

We next sought to compare the effect of soluble and particulate Ag display on B cell proliferation *in vitro*. Despite absence of co-stimulation, B cells proliferated robustly to both sHEL and pHEL equipotent stimuli when co-cultured with BAFF as assessed by vital dye dilution **(Figure 1G, H)**. As evident with 24 hr NUR77-eGFP upregulation (**Figure 1A**), proliferation in response to sHEL was highly dependent upon both dose and affinity of Ag and exhibited EC_50_ in the same order of magnitude for both assays (**Figure 1G**). By contrast, very low doses of pHELD/T (1pM *≅*1-4ng/ml HEL concentration) were sufficient to drive robust proliferation. This proceeded in a concentration-and density-dependent, but affinity-independent, manner (**Figure 1H**). Interestingly, we found that reducing the concentration of high ED SVLS allowed us to visualize a small fraction of B cells that had undergone many rounds of division **(Figure 1H, arrowheads)**. This outcome was not evident when ED was reduced nor when B cells were stimulated with sHEL **(Figure 1G, H).** These data suggest that even a small number of pHEL particles might be sufficient for maximal B cell activation; reducing pHEL concentration lowers the number of cells engaged, but not extent of activation on a single cell level, conferring an all-or-none response. Importantly, potent B cell proliferation in response to pHEL are not attributable to engagement of marginal zone (MZ) B cells – which exhibit unique biology and play a critical role in early *in vivo* responses to T-independent multivalent Ags – because these assays are performed with lymph node rather than spleen as a source of exclusively follicular B cells^44^.

### pHEL triggers highly robust BCR signaling that is insensitive to epitope affinity

We next took advantage of these unique tools to explore the biochemical basis for such robust B cell responses to Ag displayed on virus-sized particles. We first sought to dissect early signaling events downstream of BCR engagement by soluble and particulate Ag. As described above, we selected ‘equipotent’ concentrations of soluble and particulate stimuli (1pM pHEL and 1µg/ml sHEL) that induced comparable levels of downstream NUR77-GFP **(Figure 1)** in order to determine how signaling at early time points after BCR engagement differed between these modes of Ag display. We compared these stimuli to maximally potent anti-IgM Fab’2 (10µg/ml) as a reference because antibody-mediated BCR cross-linking is very well-characterized.

At early time points after stimulation (3 min), pHELD and sHELD induced comparable phospho-Erk (pErk), but signaling induced by pHELD was much more robust, peaking at 20 minutes (**Figure 2A**). Importantly, this time course was comparable whether pHEL and sHEL stimuli were pre-bound to MD4 B cells at 4C prior to 37C stimulation, or acutely admixed at 37C, implying that delayed peak was not attributable to binding kinetics of pHEL (DNS). As seen for NUR77-eGFP upregulation, naked liposomes without conjugated HEL protein failed to induce any detectable pErk. Similar to pHELD, pHELT was also highly potent for pErk induction with similar kinetics (**Figure 2A, B**). Unlike sHELD, sHELT was largely unable to trigger detectable pErk at any time point assayed (**Figure 2B, C**). Importantly, pErk levels were comparable when B cells were stimulated with either pHELT or pHELD of matched high ED (**Figure 2C**), suggesting that even early BCR signaling events do not discriminate affinity when Ag is presented on particles of viral size, and this was true across extensive titration of pHEL concentration **(Figure 2D).** S6 phosphorylation requires a lower BCR signaling threshold than Erk phosphorylation and continues to increase over the time course studied (2 hrs). Very low doses of pHELT triggered S6 phosphorylation more robustly than 1*μ*g/ml sHELT (**Supplementary Figure 2A, B**). Importantly, as seen for pErk, both pHELT and pHELD induced comparable pS6 levels and exhibited no evidence of affinity discrimination (**Supplementary Figure 2C**).

**Figure 2.**
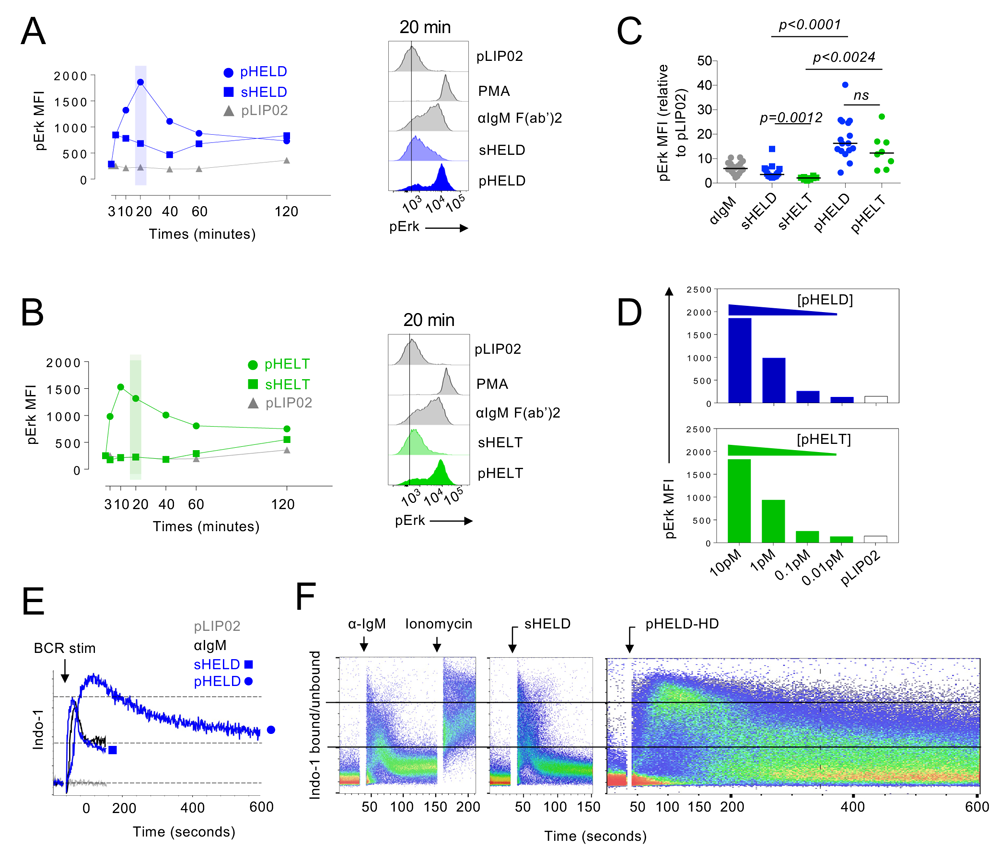
pHEL trigger robust signaling downstream of the BCR that is insensitive to epitope affinity. **A, B.** MD4 splenocytes were stimulated over a time course of 120 minutes, fixed, permeabilized, and intra-cellular pErk was measured by flow cytometry (left). Stimuli were pLIP02 = naked liposome 10pM, anti-IgM 10 μg/ml, pHELT or pHELD (256, 286 ED respectively) 1pM [liposome] ≅ 4ng/ml [HEL protein], and sHELT or sHELD 1 μg/ml. Doses of sHEL and pHEL selected were equipotent for induction of NUR77-eGFP at 24 hrs (see Fig 1C, D). Graphs (left) and histograms (right) depict pErk in B220+ cells from a single time course and is representative of 3 independent experiments. **C.** Graph depicts B cell pErk MFI after 20 min stimulation; each point represents an independent experiment and data are pooled from 18 (pHELD) or 8 (pHELT) experiments depending on stimulus. Separate groups are depicted in a single graph for convenience. Pre-specified groups were compared by a paired parametric T-test. **D.** As in A, B, but splenocytes were stimulated with decreasing concentrations of either pHELT or pHELD for 20 min. Data plotted is from a single experiment, representative of >= 4 independent experiments. **E, F.** Splenocytes and lymphocytes from MD4 mice were pooled, loaded with Indo-1 calcium indicator dye, stained to detect B220+ B cells, and stimulated as in A. Calcium entry was detected in real time by flow cytometric detection of bound and unbound Indo-1 fluorescence for 2-10 minutes. Ratio of bound/unbound fluorescence over time is plotted as population mean or for individual cells (F). Data are representative of at least 8 independent experiments.

We next measured cytosolic calcium changes following pHELD stimulation by loading MD4 B cells with the calcium indicator Indo-1 and analyzing acutely activated cells by flow cytometry. Although the response to pHELD was slower to reach peak intra-cellular calcium concentrations than with soluble stimuli (by almost 60 seconds), responding cells ultimately achieved a much higher peak that was comparable to ionomycin stimulation, suggesting it was maximal within the dynamic range of this assay **(Figure 2E, F)**. In contrast to maximal doses of either anti-IgM or sHELD which induced rapid but very short-lived intra-cellular calcium increases with uniform kinetics across the entire B cell population, doses as low as 1pM pHELD exhibited delayed but remarkably prolonged calcium increases among a subset of cells (**Figure 2E, F**). To our surprise, the response exceeded 10 minutes; B cells with intra-cellular calcium levels that were clearly above background were detectable at the latest time point assayed with only a very slow decay back to baseline. By contrast, duration of peak intra-cellular calcium triggered by soluble stimuli was on the order of 60 seconds (**Figure 2E, F**). Importantly, we do not interpret this to merely represent delayed and inhomogeneous B cell activation in response to SVLS, because nearly all B cells at the population level exhibited maximal intra-cellular calcium levels by 90 seconds following 1pM pHELD stimulation, while high intracellular calcium was still seen in a subset of cells more than 500 seconds later (**Figure 2F**). We cannot exclude the possibility that capture of additional pHEL by unoccupied BCRs does prolong calcium signaling. However, we present data later in this study to suggest that rapid signal termination in response to soluble Ag requires active engagement of inhibitory pathways that are evaded by particulate Ag.

Robust signaling triggered by pHEL was not due to selective responses by MZ B cells because lymph node samples lacking MZ B cells exhibited comparable pErk response as splenic B cells **(Supplementary Figure 2D).** Similarly, gating on follicular and MZ B cells independently on the basis of surface CD21 and CD23 expression **(Supplementary Figure 2E-I)** revealed robust signaling in both B cell subsets. Indeed, follicular B cells were reproducibly more responsive to pHEL than either immature or MZ B cells – regardless of readout. By contrast, sHEL and anti-IgM stimuli signaled most robustly in MZ B cells.

### pHEL trigger bimodal signaling that is sensitive to dose and epitope density but not affinity

Strikingly, single cell resolution of pErk responses afforded by phosflow revealed that the fraction of responding cells was reduced with lower doses of pHEL, but even very low concentrations of pHELT or pHELD could induce comparable and maximal levels of pErk in a small fraction of B cells in culture (**Figure 3A**). This bimodal signaling was reminiscent of the proliferative response of B cells to pHELT/D with high ED (**Figure 1H**). Reducing ED also reduced pErk MFI but in a bimodal manner, suggesting “all-or-none” digital responses to pHEL by single B cells (**Figure 3B**). This observation extended as well to pHEL with ultralow ED (pHELT ED 3). Strikingly, such ultralow ED particles were nevertheless many orders of magnitude more potent than soluble Ag, inducing robust pErk response at 10pM SVLS (equivalent to 30pM *≅* 0.4ng HELT protein) (**Supplementary Figure 2J**). This corresponded well to robust Nur77-eGFP induction by these reagents (**Supplementary Figure 1F**) and reinforced the possibility that pHEL potency was not merely attributable to extensive BCR cross-linking by individual SVLS particles.

**Figure 3.**
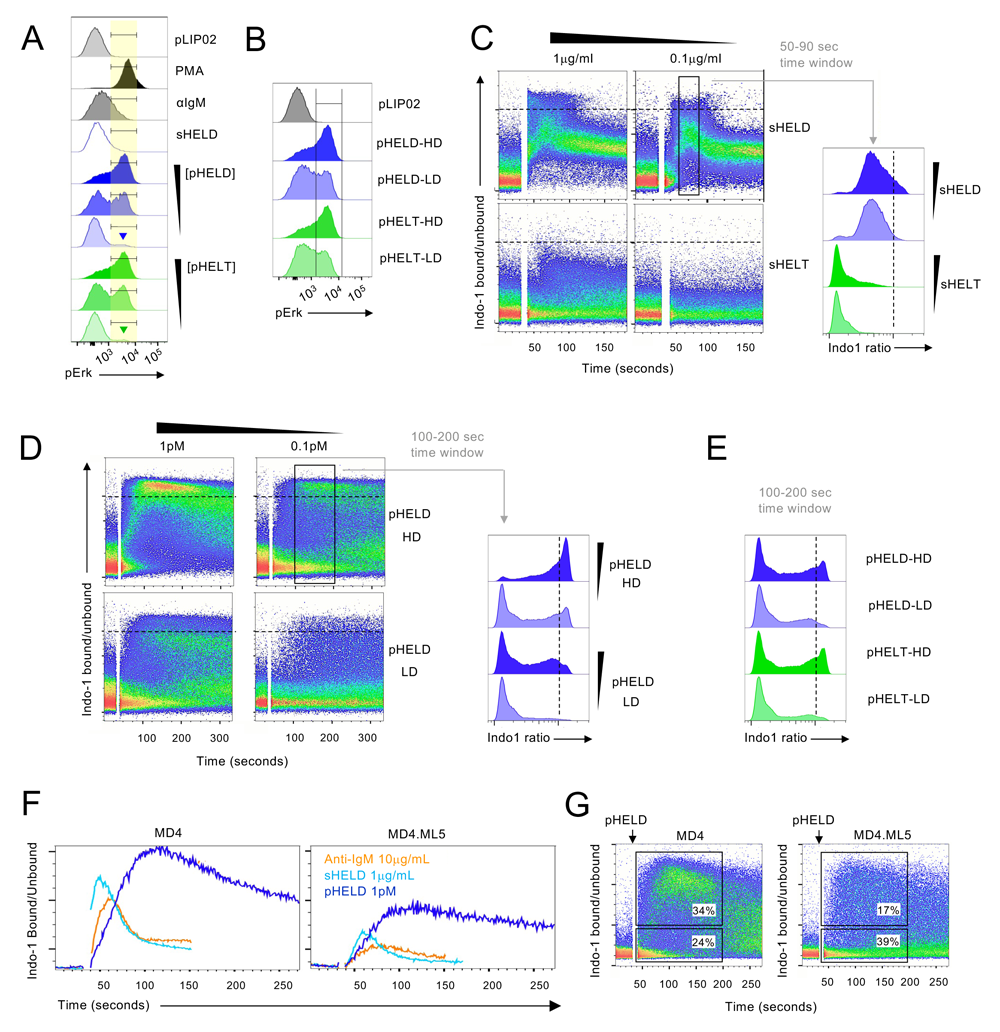
Particulate Ag trigger bimodal signaling that is sensitive to dose and epitope density, but not to epitope affinity. **A.** As in 2A/B, but splenocytes were stimulated with decreasing concentrations of either pHELT or pHELD (10pM, 1pM, 0.1pM). Samples correspond to MFI quantification in 2D. Inset goal posts gate on subset of cells with maximal pErk signal. **B.** As in A, but splenocytes were stimulated with 10pM of SVLS carrying either high density (HD) or low density (LD) of HELD or HELT (286, 256, 61, 64 respective ED). Data in A, B is representative of at least 4 independent experiments. **C-E.** Pooled splenocytes and lymphocytes from MD4 mice were loaded with Indo-1 calcium indicator dye, stained with surface markers to detect B220+ cells and in C stimulated with sHELD/T (1 and 0.1 μg/ml). In D cells are stimulated with 1pM or 0.1pM of high or low density pHELD (286, 23 ED respectively), and in E are stimulated with high or low density pHELD/T (286/256, 61/64 ED respectively). Calcium entry was assessed by flow cytometry for indicated time. Plots depict geometric mean of Indo1 bound/unbound fluorescence ratio in individual cells over time (seconds) on a log scale. Histograms display Indo1 bound/unbound fluorescence ratio gated on a defined time window to capture peak cytosolic calcium for each type of stimulus. Data in C-E depict same Indo-1 axis/scale and dashed black line marks corresponding value across dot plots and histograms for comparison. Data are representative of at least 3 independent experiments depending on stimuli. **F, G**. As in 2E, F and 3C except cells from MD4 or MD4.ML5 mice were stimulated with indicated reagents: anti-IgM 10 μg/ml, sHELD 1 μg/ml, or pHELD-HD (ED 286) 1pM. Indo-1 ratio is shown on a linear scale. Gated in G are corresponding time gates to capture fraction of responding or non-responding B cells. Data are representative of 2 independent experiments.

To our surprise even at the earliest time points we could assess, all pHEL calcium responses were bimodal, in marked contrast to soluble stimuli (**Figure 3C-E**); as with pErk, reducing concentration of pHEL decreased fraction of responding cells but not intra-cellular calcium level among responding cells (**Figure 3A, D**). These bimodal calcium responses were sensitive to dose and ED, and strikingly appeared to give a reduced maximal Indo-1 (ca2+ bound/unbound) ratio with low ED. Importantly, these responses to SVLS were insensitive to affinity in marked contrast to soluble stimuli. Such bimodal single cell behavior in response to pHEL suggests that particulate Ag triggers signal amplification that may operate at a node upstream of both MAPK and calcium pathways.

We next took advantage of a well-studied model of B cell tolerance; Hy10 BCR Tg B cells that develop and mature in hosts expressing sHEL as a self-antigen (MD4.ML5 Double Tg mice) exhibit canonical features of anergy, including IgM downregulation and suppressed proximal BCR signal transduction^42, 45^. We found that both soluble stimuli and pHEL triggered dampened responses in self-reactive, anergic MD4.ML5 B cells relative to naïve MD4 B cells (**Figure 3F**). However, calcium increases triggered by 1pM pHEL in anergic B cells was nevertheless more robust and considerably more prolonged than maximal soluble stimulation of naïve MD4 B cells (**Figure 3F**). Perhaps most strikingly, pHEL triggered bimodal behavior; a fraction of anergic MD4.ML5 cells exhibited maximal calcium entry that was comparably robust and prolonged relative to responding non-anergic, naive MD4 B cells (**Figure 3G**). This suggested that a fraction of even deeply anergic B cells could be “re-awakened” by virus-like Ag display. Indeed, ncreased amplitude and duration of cytosolic calcium increases can in turn activate transcriptional pathways such as NF-*κ*B that are normally suppressed by anergy^46, 47^.

### pHEL requires canonical BCR signaling pathway but is independent of MYD88 and IRAK1/4

Because pHEL trigger remarkably robust signal transduction in B cells, we sought to exclude a role for any contaminating nucleic acids that could be delivered via pHEL to endosomal TLRs. To do so, we first took advantage of a highly selective IRAK1/4 inhibitor (R568, Rigel Pharmaceuticals). This drug blocked CpG-induced pErk and pS6 but did not inhibit pHEL signaling **(Figures 4A, B)**. Similarly, deletion of MYD88 completely suppressed signaling by LPS and CpG but had no impact on other pathways (such as BCR or CD40) as expected (**Figure 4C, D**). Importantly, MYD88 was completely dispensable for pHEL responses *in vitro* (**Figures 4C, D**). Conversely, pHEL responses were fully dependent upon activity of canonical BCR pathway signaling molecules like SYK, BTK and PI3K^48^; small molecule inhibitors of these enzymes completely eliminated downstream induction of pErk in response to pHEL, but did not impair CpG-induced Erk phosphorylation via endosomal TLR9 (**Figure 4E-H)**.

**Figure 4.**
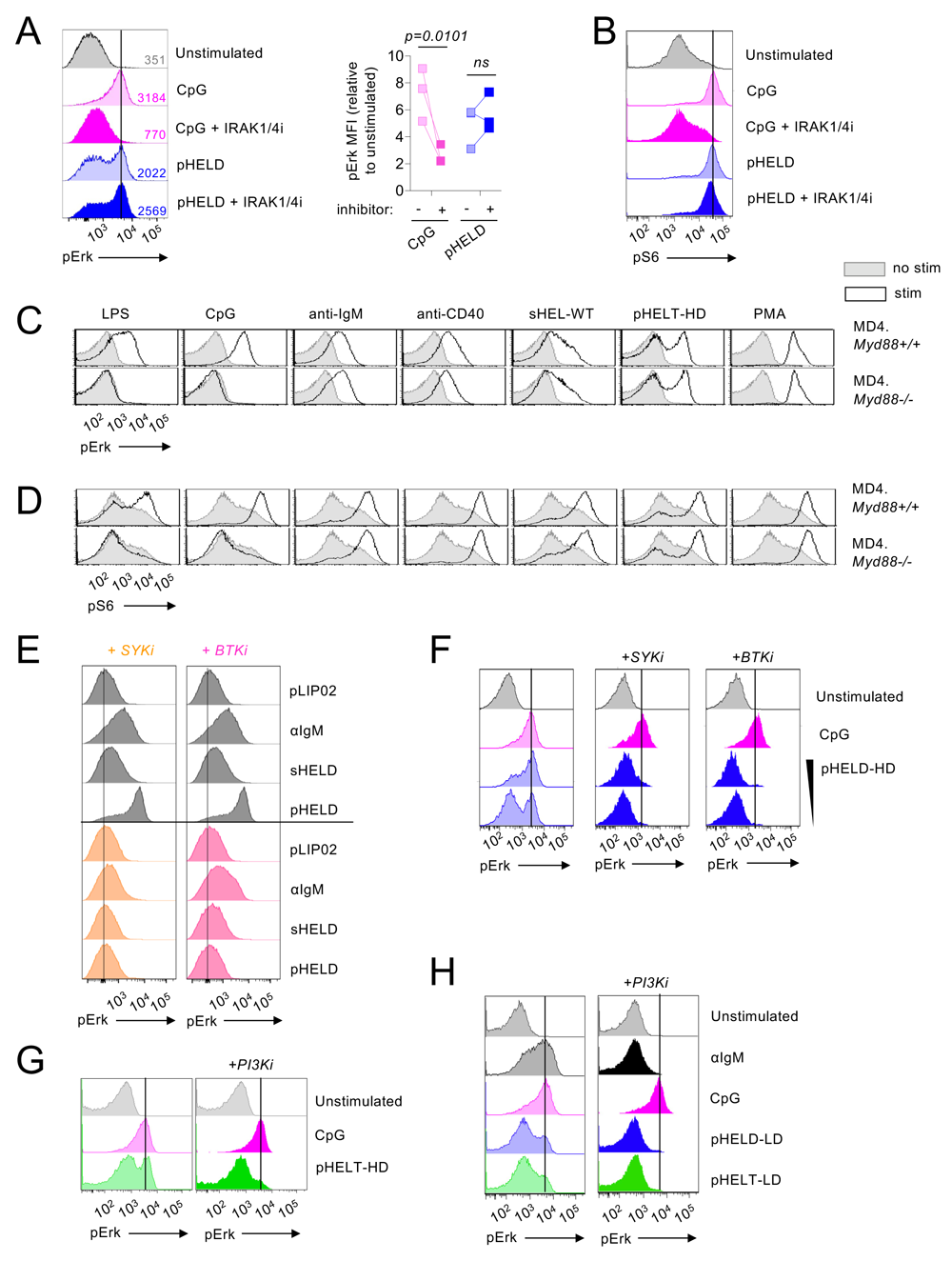
pHEL signaling is independent of MYD88/IRAK1/2 but requires canonical BCR pathway. **A, B.** Pooled splenocytes and lymphocytes from MD4 mice were pre-incubated at 37C for 15 min with Rigel R568 IRAK1/4 inhibitor (1 μM) followed by 20 min incubation with indicated stimuli (2.5 μM CpG, 1pM pHELD) and then fixed, permeabilized, and stained to detect pErk or pS6 as well as B220. Representative histograms depict pErk (A) or pS6 (B) in B220+ B cells. Graph depicts pErk MFI from 3 independent experiments. Data in B is representative of at least 4 independent experiments. **C, D.** Pooled splenocytes and lymphocytes from MD4 *Myd88*+/+ and MD4 *Myd88*–/– mice were stimulated with indicated stimuli for 20 minutes and stained to detect pErk or pS6 in B220+ B cells as in A, B (LPS 10 μg/ml, CpG 2.5 μM, anti-IgM 10 μg/ml, anti-CD40 1 μg/ml, sHEL-WT 1 μg/ml, or pHELT-HD (ED 256) 1pM,). Histograms depict pErk (C) or pS6 (D) and are representative of 4 (C) or 3 (D) independent experiments. **E, F.** As in 4A except pre-treatment with SYKi (Bay 61-3606 1 μM) or BTKi (ibrutinib 100mM), followed by 20 minute incubation with indicated stimuli: pHELD-HD (ED 286) 1pM and 0.1pM (F), sHELD 1 μg/ml, anti-IgM 10 μg/ml, 2.5 μM CpG. Histograms are representative of 4 independent experiments. **G, H.** As in 4A except pre-treatment with PI3Ki (Ly290049 10 μM) and stimuli (pHELT-HD ED=256 1pM in G; pHELD-LD ED=53 and pHELT-LD ED=58 1pM in H). Histograms are representative of at least 4 independent experiments.

### pHEL trigger robust signal amplification downstream of the BCR

Robust bimodal or digital signaling responses suggest that pHEL may trigger signal amplification downstream of the BCR relative to soluble Ag. To investigate this mechanism, we comprehensively analysed the most proximal biochemical events triggered in B cells at early time points after stimulation with anti-IgM, sHELD and pHELD. We selected a 3-minute stimulation for Western blot analysis, because pHELD and sHELD induce comparable Erk phosphorylation at this time point as detected by intracellular staining (**Figure 2A**). Although downstream readouts like pErk and pIKK were robustly induced by both sHEL and pHEL after 3 minutes, to our surprise total tyrosine phosphorylation (probed with 4G10 mAb) was virtually undetectable following pHEL stimulation (**Figure 5A, B**). Similarly, tyrosine phosphorylation of the BCR itself (CD79A), SYK, and PI3K were also robustly induced by soluble stimuli, but virtually undetectable by western blot for pHEL stimulation **(Figure 5C, D)**. Phosphorylation of PLC*γ*2 at Y1217 is a key proximal event triggered by BCR engagement that occurs upstream of BTK^49^. This was also induced by soluble stimuli, but not increased with pHEL. Nevertheless, although phosphorylation of proximal signaling nodes in response to pHEL was below the limits of detection by western blotting, pHEL responses do engage and require enzymatic activity of canonical BCR signaling molecules SYK, BTK, and PI3K because small molecule inhibitors of these enzymes completely prevented downstream induction of pErk in response to pHEL as noted earlier **(Figure 4E-H)**^48^.

**Figure 5.**
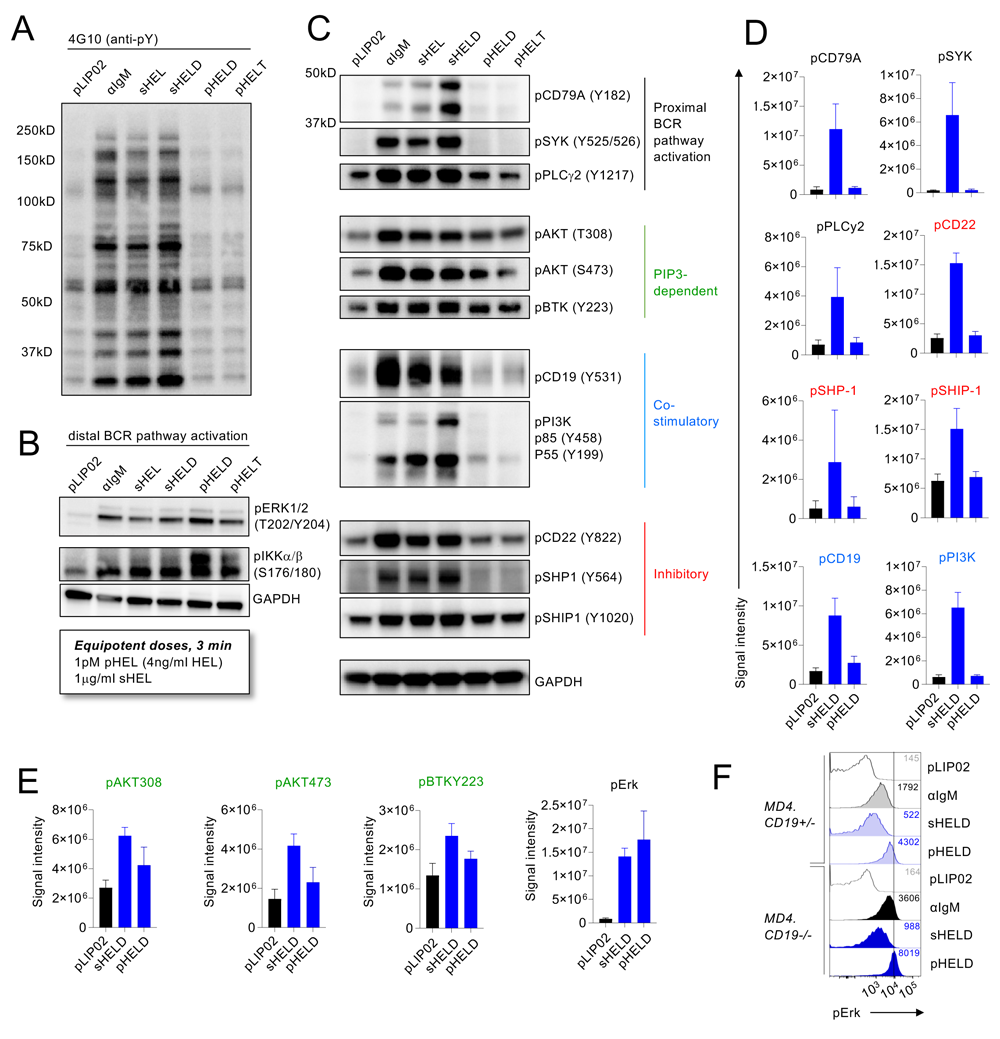
pHEL trigger robust signal amplification downstream of the BCR. **A-C.** MD4 splenocytes and lymphocytes were pooled, and B cells were isolated by bench-top negative selection. B cells were stimulated for 3 min at 37C with either naked liposome (pLIP02 1pM), 10 μg/ml anti-IgM, 1 μg/ml sHEL-WT / sHELD, or 1pM pHELD / pHELT (high ED 250-290). Lysates were analysed by western blotting to detect total phospho-tyrosine (4G10) or specific phospho-species in the BCR signaling cascade. GAPDH (panel B and C, bottom) is shown as a loading control because a common pool of cells was used for all stimulation conditions. Images depict blots performed with a single set of lysates representative of 3-6 independent experiments depending on blotting Ab with corresponding quantification across biological replicates in D, E. **D, E.** Summary data of phosphoprotein quantification from 3-4 independent experiments. Quantification shows raw band intensity without normalization. Rather, in each experiment, blots were probed for GAPDH as depicted in B to confirm accurate loading. Error bars depict SEM. **F.** 20’ Phosflow assay to detect pErk in *Cd19*+/– or *Cd19*–/– MD4 splenocytes after stimulation with either control pLIP02 1pM, 10 μg/ml anti-IgM, 1 μg/ml sHELD, or 1pM pHELD-HD. Samples were co-stained with B220, CD23, and CD21 to detect B cell subsets. Histograms depict CD23+ B cells and are representative of 4 independent experiments.

Consistent with calcium kinetics (**Figures 2E, F, 3D**), robust pErk and pIKK induction by pHEL indicated that BCR engagement occurred by 3 minutes (**Figure 5B**) and that undetectable phosphorylation of the proximal BCR signaling cascade was not attributable to selection of an early time point. This discordance between proximal and distal signaling led us to hypothesize that a small number of BCRs engaged by low concentration pHEL induced proximal tyrosine phosphorylation below the limits of detection, and subsequent downstream signal amplification resulted in robust downstream phosphorylation of Erk and IKK. Conversely, these data suggest that potent downstream signaling by pHEL was not merely a consequence of enhanced proximal BCR engagement (i.e. extensive BCR cross-linking) as such a mechanism would be anticipated to produce robust phosphorylation at the most proximal signaling nodes such as pCD79A and pSYK. This argues against mere avidity alone as a mechanism for pHEL potency.

The second messenger PIP3 represents a key signaling intermediate that dynamically accumulates at the plasma membrane to nucleate the assembly of the BCR ‘signalosome’ and serves as a node for tuning BCR signal amplification^50^. AKT T308 phosphorylation is mediated by PDK1 and is highly dependent upon PIP3 for recruitment of both AKT and PDK1 via their PH-binding domains^51, 52^. By contrast to proximal tyrosine phosphorylation, detectable AKT serine/threonine phosphorylation was induced by both sHEL and pHEL at 3 minutes (**Figures 5C-E**). BTK autophosphorylation at Y223 also depends upon recruitment to PIP3 and dimerization^53, 54^, and is measurably induced by pHEL (**Figure 5C, E**). Detectable phosphorylation of these sites following pHEL stimulation suggested that local PIP3 generation by PI3K at the membrane might serve to disproportionately amplify signaling in response to pHEL. Importantly, although PI3K phosphorylation is undetectable after pHEL stimulation (**Figures 5C, D**), downstream pHEL signaling is nevertheless sensitive to PI3K inhibition (**Figure 4G, H**).

### Potent signaling by pHEL *in vitro* is not dependent upon CD19

PIP3 abundance at the membrane in response to BCR stimulation is regulated both at the level of its generation and its degradation^48, 50^. The CD19 coreceptor plays an important role in amplifying PIP3 production by PI3K in response to BCR stimulation. Moreover, CD19 is co-clustered with BCR upon encounter with cell–membrane associated Ags or via complement-coated Ags, and CD19 is required for highly robust responses to such Ag, but it is dispensable for *in vitro* response to soluble Ag^25, 27, 29, 55^. CD19 phosphorylation was robustly induced by anti-IgM and sHEL, but not by pHEL, raising the possibility that CD19 was not engaged by pHEL (**Figure 5C, D**). We therefore generated CD19-deficient MD4 mice to test whether CD19 engagement could account for potent pHEL responses. In contrast to signaling triggered by Ag displayed on a cell membrane or via actin depolymerization^27, 29^, CD19 is dispensable for robust signaling by particulate Ag *in vitro* **(Figure 5F)**.

### pHEL trigger robust signal transduction downstream of the BCR by evading LYN-dependent inhibitory pathways

PIP3 abundance at the membrane and proximal BCR signaling are both negatively regulated by ITIM domain-containing co-receptors and associated inhibitory phosphatases^48, 50^. We observed robust phosphorylation of CD22, pSHP1, pSHIP1 in response to anti-IgM and sHEL, but not pHEL (**Figure 5C, D**). This raised the possibility that particulate Ag circumvents engagement of inhibitory coreceptors to drive potent B cell responses. Indeed, exclusion of inhibitory coreceptors and associated phosphatases is postulated to contribute to potent responses by cell membrane-associated Ags^15, 20, 25, 26^. Several Src family kinases (SFK) are expressed in B cells and can all mediate ITAM-dependent signaling downstream of BCR engagement. By contrast, the SFK LYN plays a non-redundant role in phosphorylation of ITIM domains of inhibitory co-receptors; LYN-deficient B cells therefore fail to engage downstream phosphatases SHP1 and SHIP1 in response to anti-IgM stimulation, but can activate ITAM-dependent BCR signaling (as well as CD19) via other SFKs (**Figure 6A**)^48, 50, 56, 57^. To test the hypothesis that pHEL evades engagement of inhibitory co-receptors to trigger highly potent responses – in contrast to soluble stimuli – we generated MD4 B cells deficient for LYN to selectively eliminate all ITIM-dependent signaling. In the absence of LYN, anti-IgM and sHEL-induced pErk and calcium entry were markedly amplified (**Figure 6B-D**). Even sHELT which triggers virtually undetectable pErk in LYN-sufficient B cells exhibit highly enhanced signaling comparable to pHELT (**Supplementary Figure 3A**). By contrast, pHEL responses were not enhanced in MD4.Lyn–/– B cells; no amplification of pHEL signals irrespective of affinity, ED, or dose could be detected in the absence of LYN – even for SVLS with ultralow ED (**Figure 6B-D, Supplementary Figure 3A, B)**. As a result, in MD4.*Lyn*–/– B cells, sHELD and pHELD induced comparably robust pErk and calcium entry **(Figure 6B-D)**, indicating that the evasion of LYN-dependent inhibitory tone by pHEL but not sHEL accounts, at least in part, for the remarkable potency of pHEL in LYN-sufficient B cells. Strikingly, pHEL-induced calcium in MD4 B cells phenocopy the heightened, delayed and prolonged kinetics of calcium responses by LYN-deficient cells to soluble stimuli (**Figure 6D**). Conversely, not only amplitude, but also kinetics of pHEL-induced calcium entry was unaltered by LYN-deficiency (**Figure 6D**). This data further suggests that curtailed signaling triggered by soluble stimuli relative to pHEL was not simply attributable to saturating BCR occupancy and/or internalization of surface receptor as this would not be rescued by LYN-deficiency. Rather we conclude that both signal amplitude and signal duration in response to soluble – but not particulate – Ag is constrained by ITIM engagement.

**Figure 6.**
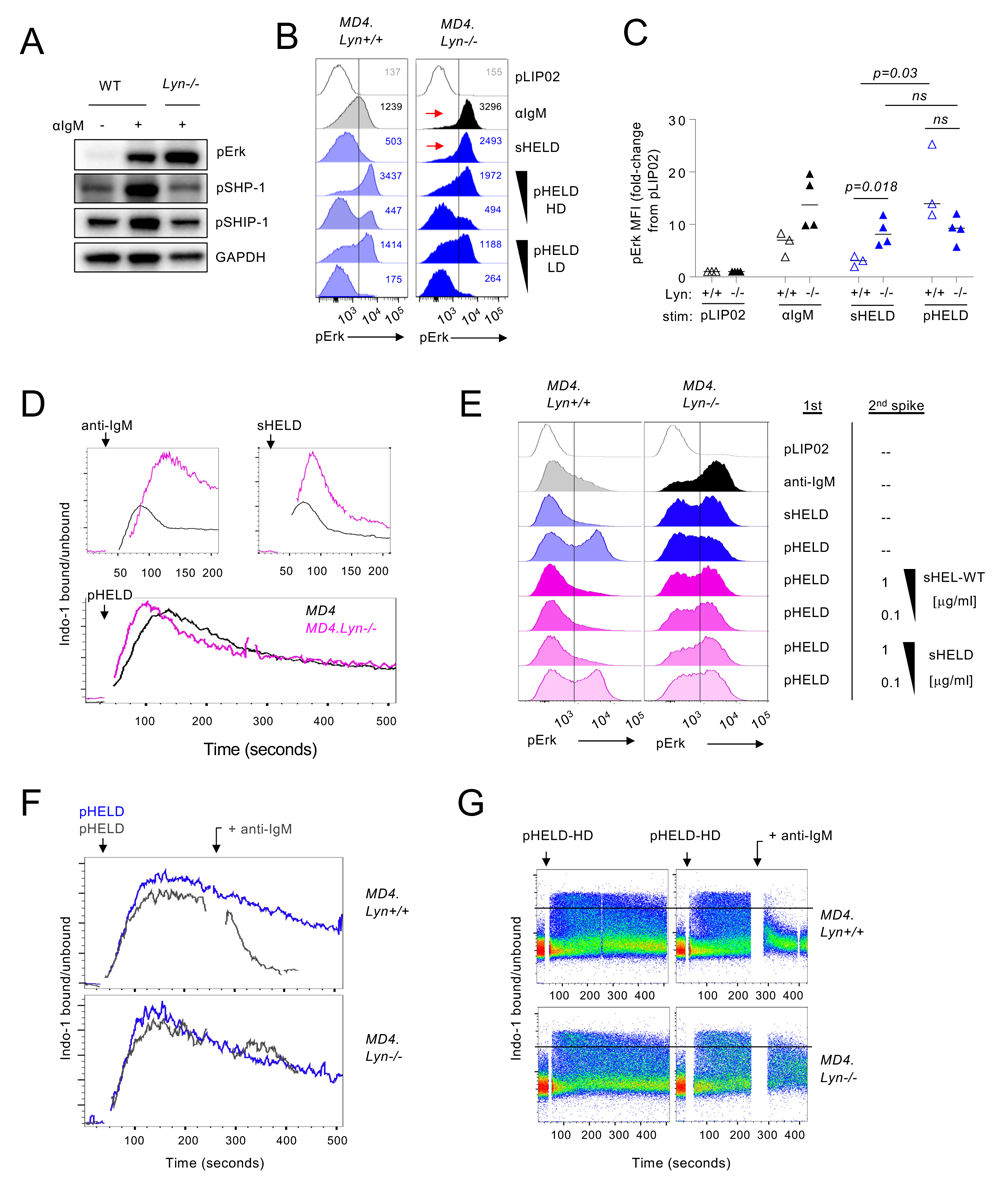
Soluble but not particulate Ag are restrained by LYN-dependent inhibitory signaling. **A.** Lysates were generated and probed as in 5A-C, except either WT or *Lyn*–/– polyclonal B cells were stimulated for 3 minutes with 10 μg/ml anti-IgM. Blot is representative of at least 3 independent experiments. **B.** 20’ Phosflow assay to detect pErk in *Lyn*+/+ and *Lyn*–/– MD4 pooled splenocytes and lymphocytes after stimulation with either control pLIP02 1pM, 10 μg/ml anti-IgM, 1 μg/ml sHELD, or pHELD. Both high density and low density (HD/LD 286, 61) pHELD were used at two doses each (1, 0.1pM). Samples were co-stained with B220, CD23, and CD21 to detect B cell subsets. Inset values represent MFI. Histograms depict CD23+ B cells and are representative of at least 8 independent experiments. **C.** Graph depicts data in B pooled from 3-4 independent experiments. Pre-specified groups were compared by parametric T-tests and are plotted together on the same graph for convenience. **D.** Pooled splenocytes and lymphocytes from *Lyn*+/+ and *Lyn*–/– MD4 mice were loaded with Indo-1 calcium indicator dye, stained with CD21 and CD23 surface markers to detect B220+ subsets, and stimulated with anti-IgM (10 μg/ml), sHELD (1 μg/ml), or pHELD-HD (ED 289, 1pM). Calcium entry was assessed by flow cytometry for indicated time as in 2E, F. Plots depict geometric mean of Indo1 bound/unbound fluorescence ratio over time in CD23+ B cells (seconds). Data are representative of 3 independent experiments. **E.** Splenocytes from *Lyn*+/+ and *Lyn*–/– MD4 mice were incubated with 1^st^ stimulus at 4C for 15 minutes, followed by 2^nd^ add-in stimulus (or media) for another 15 min at 4C. Cells were then incubated for 20’ at 37C, and fixed, permeabilized, and stained as in 2A to detect pErk. 1^st^ stimuli: 10 μg/ml anti-IgM, 1 μg/ml sHELD, or 5pM pHELD-HD (ED 289). 2^nd^ add-in doses of sHEL-WT or sHELD as annotated (μg/ml). histograms depict B220+ cells and are representative of 4 independent experiments. **F, G.** As in D except with late addition of 10μg/ml anti-IgM following pHEL stimulation at indicated time (gray kinetic curve). Data in F depict geometric mean of Indo1 bound/unbound fluorescence ratio over time in CD23+ B cells, while F depicts same samples with single cell reslution. Data are representative of at least 3 independent experiments.

### Soluble stimuli dominantly suppress responses to pHEL by engagement of LYN-dependent inhibitory pathways

Our data suggests that LYN-dependent inhibitory pathways are engaged and restrain responses to soluble stimuli such as sHEL and anti-IgM, but are evaded by pHEL. We next sought to test whether enforced engagement of LYN-dependent inhibitory pathways by soluble stimuli is sufficient to suppress pHEL signaling. To do so, we developed an assay in which pHEL were pre-bound to MD4 B cells at 4C to forestall signaling, followed by subsequent introduction of soluble stimuli. Cells were then assessed for pErk induction after 20 min incubation at 37C to permit signal transduction. The rationale for this approach is to first ensure BCR engagement by high ED pHEL (which is expected to bind with high avidity). Subsequent introduction of soluble stimuli bound to abundant free BCRs are predicted to dominantly co-engage inhibitory pathways. Importantly, kinetics of signaling by pHEL are not altered by this incubation strategy and still peak at 20 minutes in the absence of sHELD (**Figure 6E** and DNS). By contrast, treatment of pHEL-bound cells with sHEL was sufficient to suppress signaling and this suppression required LYN (**Figure 6E, Supplementary Figure 3C**). Such suppression by sHEL was both affinity-and dose-dependent. Importantly, this suppression was not merely due to competition for receptor binding by sHEL-WT as lower affinity sHELD could also suppress signaling by pre-bound high ED pHELD (**Figure 6E**), as could anti-IgM treatment (which neither engages IgD BCRs nor competes with HEL epitope for binding of the Hy10 IgM BCR) (**Supplementary Figure 3C**). In all cases, such suppression was rescued in the absence of LYN (**Figure 6E, Supplementary Figure 3C**).

These data suggest that co-ligation with soluble stimuli can prevent signaling by pHEL in a LYN-dependent manner. We next sought to determine whether such soluble stimuli, via LYN engagement could terminate signaling *after* initiation. To do so, we adapted a flow-based calcium assay to trigger cytosolic calcium increases with pHELD and subsequently add either soluble stimulus (or control) once peak of the response is reached (**Figure 6F, G, Supplementary Figure 3D**). Indeed, following pHEL stimulation cytosolic calcium remains at maximal levels in a fraction of WT MD4 B cells and this signal is rapidly terminated with addition of soluble stimuli; we found that addition of either anti-IgM or sHELD was sufficient to completely suppress prolonged pHEL calcium responses. Importantly, this was completely LYN-dependent in the case of anti-IgM and partially LYN-dependent for sHELD. These data suggest that signal termination is neither due to displacement of pHEL from BCRs, nor to internalization of BCRs because anti-IgM treatment neither blocks nor removes IgD BCRs from the B cell surface. Rather, we conclude that dominant engagement of LYN-dependent inhibitory signaling by soluble stimuli suppresses B cell responses to particulate Ag.

### pHEL induces a highly robust NF-κB signal that mimics CD40 stimulation

When comparing differences in signal transduction induced by sHEL and pHEL, we noted that NF-κB activation was disproportionately enhanced by pHEL (**Figure 4A, Figure 7A**). BCR activation elicited by sHEL or anti-IgM cross-linking only resulted in weak phosphorylation of IKK*β*, the catalytic subunit required for its kinase activity and downstream NF-κB translocation to the nucleus^58^. However, IKK*β* was highly phosphorylated in response to pHEL **(Figure 7A)** and correspondingly, IκB*α* (the substrate of IKK catalytic activity) was rapidly and robustly degraded in response to pHEL but not soluble stimuli like sHEL and anti-IgM **(Figure 7B-D)**. As with other signaling nodes interrogated, NF-κB activation was independent of pHEL Ag affinity but sensitive to pHEL ED and concentration **(Figure 7B, E)**. Since the degradation of IκB*α* is required to release p65 to translocate to the nucleus, we isolated nuclei from stimulated B cells and directly probed for p65 in permeabilized nuclei. In agreement with IKK phosphorylation and IκB*α* kinetics, we show that p65 is found in high abundance in the nucleus of B cells stimulated with pHEL **(Figure 7F)**.

**Figure 7.**
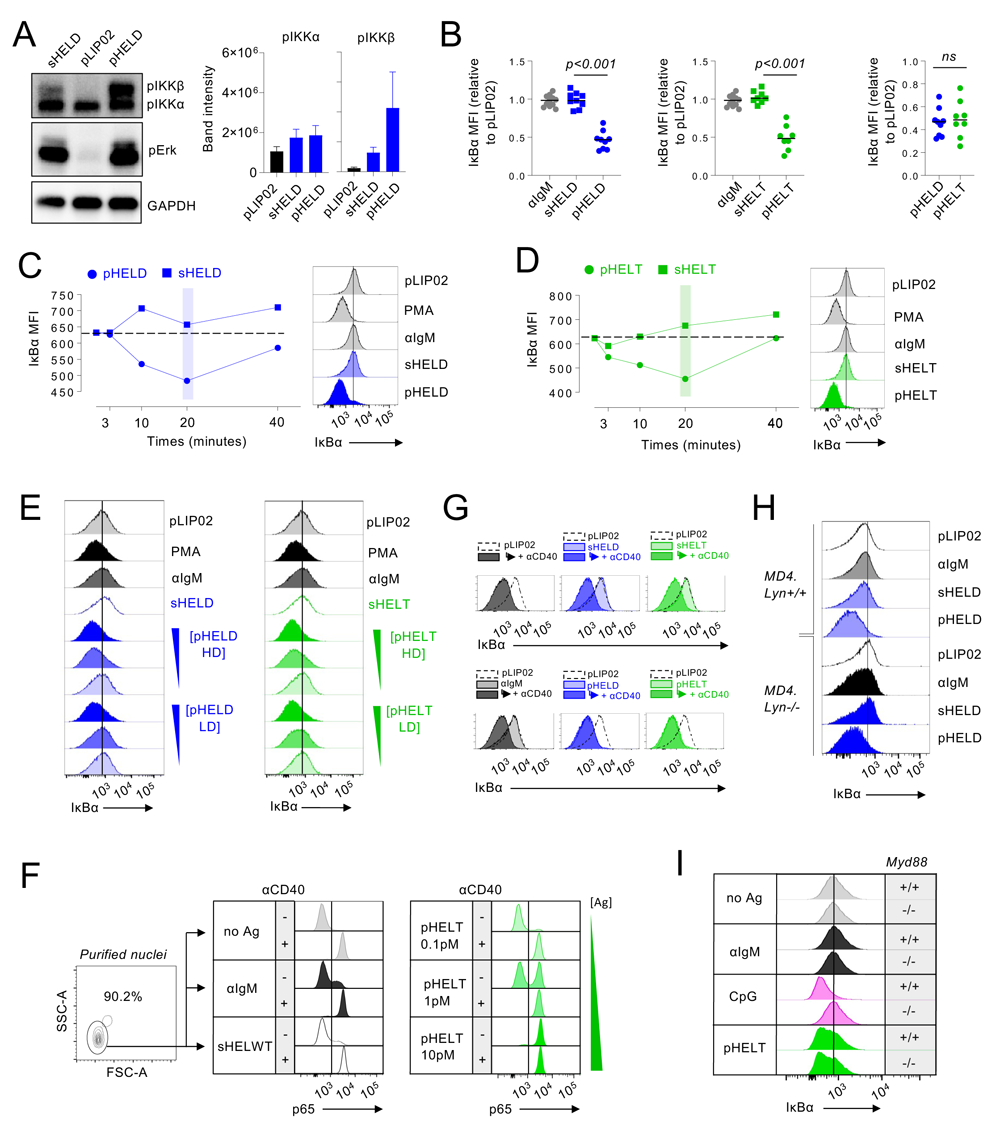
pHEL trigger highly robust NF-κB signaling that mimics co-stimulation and is independent of MYD88. **A.** Lysates were prepared as in Fig 5 but probed for pIKKα/β and pErk as shown in 5B. Western blot images are representative of 4 independent experiments. Quantification (right) show raw band intensity for IKKα and IKKβ subunits, error bars show SEM. **B-D.** As in Fig 2A/B, pooled splenocytes and lymphocytes from MD4 mice were stimulated over a time course, fixed, permeabilized, and stained to detect total intra-cellular IκBα by flow cytometry. Graph (B) depicts IκB MFI in B220+ cells at 20 minutes. Each data point represents an individual experiment pooled from >=8 independent experiments. Groups were compared by a parametric T-test. Stimulation with either sHELD/pHELD-HD (C) or sHELT/pHELT-HD (D) is shown across a time course (left) and representative histograms (right) depict IκB degradation in B220+ cells after 20 minutes. **E.** As in B-D except histograms showing IκBα degradation at 20 minutes post-stimulation for soluble stimuli sHELD/T and pHELT/D with high or low ED (HD vs. LD). Inset triangles represent decreasing concentration of pHEL (10, 1, 0.1pM). Data are representative of >= 4 independent experiments. **F.** B cells were isolated by negative selection from MD4 spleen and lymph nodes. Cells were stimulated for 20 minutes (10ug/ml anti-IgM, 0.1, 1, 10pM pHELT-HD, 1μg/ml sHEL-WT +/- 1 μg/ml anti-CD40), and nuclei were purified by centrifugation against a density gradient. Nuclei were then permeabilized and stained. Nuclei were gated based on FSC and SSC as shown in representative plot (left) and interrogated for nuclear translocated p65 (right). Data are representative of 3 independent experiments. **G.** Purified B cells were stimulated in the presence or absence of anti-CD40 (1 μg/ml) as well as sHEL 1 μg/ml or pHEL 1pM, and IκBα degradation was measured by intracellular staining 20 minutes later. Each overlaid histogram shows internal negative control pLIP02 as reference (dotted histogram). Light shaded histograms show BCR stimulus condition alone, dark shaded histograms show BCR stimulus condition + anti-CD40. Data are representative of >= 3 independent experiments. **H.** As in 6B, except splenocytes from *Lyn*+/+ and *Lyn*–/– MD4 mice were stained to detect intracellular IκB in CD23+ B cells at 20 minutes. Data are representative of 3 independent experiments. **I.** As in 4C, D except stained to detect intracellular IκB in CD23+ B cells at 20 minutes. Data are representative of 3 independent experiments. In panels C, D, E, H, I: line in offset histograms references internal negative control.

It has been proposed that mature naïve follicular B cells require two signals in order to mount an immune response: signal 1 delivered by the BCR and signal 2 delivered by T cell help, in part via CD40L-CD40 interaction. CD40 is a TNF-receptor superfamily member which strongly activates NF-κB pathway. By contrast, soluble Ag and even anti-IgM cannot trigger comparably robust NF-κB activation at any dose *in vitro* and this may help confer dependence on signal 2. Since pHEL readily drives highly robust NF-κB activation relative to soluble Ag, we hypothesized that Ag displayed on particles of viral size might mimic co-stimulation and bypass the requirement for T cell help. Indeed, immunization with pHEL is sufficient to produce Ab response in the absence of T cell help and without adjuvant (biorxiv preprint 2023)^59^. We therefore stimulated B cells with either sHEL or pHEL in the presence or absence of CD40 ligation. As expected, CD40 stimulation triggered robust IκB*α* degradation and nuclear translocation of p65 both independently and in combination with soluble BCR stimuli **(Figure 7F, G)**. However, CD40 stimulation produced no additional increase in NF-κB activation when combined with pHEL. Rather, pHEL alone drives IκB*α* degradation and p65 nuclear accumulation that is comparable in magnitude to that induced by CD40 cross-linking, suggesting that BCR activation by pHEL was sufficient to convey costimulatory-like information. Moreover, pHEL exhibited highly bimodal p65 nuclear translocation in a dose-dependent manner such that even low doses of pHEL induced maximal signal in a subset of cells (**Figure 7F**). Interestingly, unlike for pErk and calcium entry **(Figure 6B-D)**, the genetic absence of LYN was not capable of rescuing the defect in IκB degradation by sHEL, but did so partially in response to anti-IgM (**Figure 7H)**. This observation implies that capacity of pHEL to trigger robust NF-κB activation is not completely accounted for by evasion of inhibitory co-receptor engagement, and that other unique features of dense Ag display on virus-sized particles efficiently link BCR to NF-κB activation. Importantly, IκB*α* degradation by SVLS was independent of MYD88 (**Figure 7I, Supplementary Figure 3E)**.

### pHEL drive very robust B cell growth, MYC induction, and survival in the absence of co-stimulation

We next sought to explore the functional consequences of potent, prolonged, and qualitatively distinct signaling by particulate Ag display. We analyzed downstream cellular outcomes including cell growth, survival, and proliferation. We found that cell growth – as measured by forward scatter by flow cytometry – was greatly enhanced in response to pHEL relative to sHEL after 24 hours **(Figure 8A)**. These data suggested that cell growth induced by particulate Ag could exceed that triggered by maximal soluble Ag stimulation. Because cell growth is associated with cMYC upregulation^60, 61^, we assayed MYC induction in response to both soluble and particulate stimuli. After 24 hrs, we found that – relative to other markers of B cell activation, including CD69 and CD86 – 1pM pHELT induced higher intra-cellular MYC protein expression than maximally potent doses of sHEL-WT and anti-IgM (**Figure 8B)**. This suggested that qualitatively distinct signaling by pHEL efficiently promotes and sustains MYC protein accumulation. Consistent with this observation, we also found that pHEL promotes cMYC expression disproportionately to NUR77-eGFP upregulation after 24 hr stimulation (**Supplementary Figure 4A).** Consistent with robust NF-*κ*B activation and MYC induction, at successive timepoints in culture we found that B cell survival was consistently enhanced by pHEL relative to sHEL even with addition of the B cell survival factor BAFF **(Figure 8C, Supplementary Figure 4B)**. Importantly, neither MYC nor CD69 nor cell growth triggered by pHEL was dependent on MYD88 (**Figure 8D, E, Supplementary Figure 4C**).

**Figure 8.**
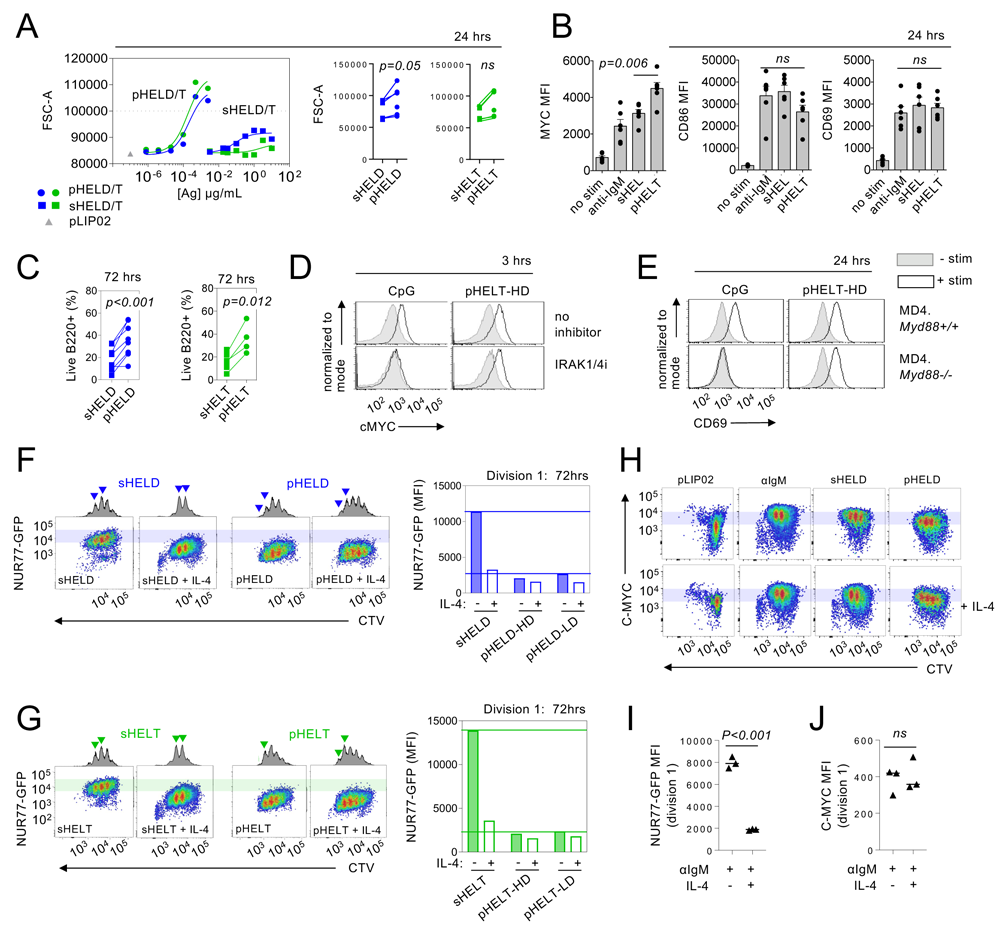
pHEL trigger robust proliferation with a lower NUR77-eGFP threshold than soluble stimuli. **A.** Pooled splenocytes and lymphocytes from MD4 mice were incubated with indicated stimuli for 24 hours, stained to detect B220+ cells and analysed by flow cytometry. Graphs depicting dose-response curve for FSC-A MFI (A) are representative of at least 4 independent experiments. Summary data correspond to 24 hr stimulation and show FSC-A values for highest concentration of sHELD/T and pHELD/T-HD in each individual experiment (1 μg/ml and 1pM respectively) for N=5-6 experiments. Lines connect samples from the same experiment. Groups were compared by a parametric paired T-test. **B.** Pooled splenocytes and lymphocytes from MD4 mice were incubated with indicated stimuli for 24 hours (anti-IgM 10 μg/ml, sHEL-WT 1 μg/ml, and pHELT-HD 1pM), and stained to detect B220 and either intra-cellular MYC or surface CD69 and CD86 expression by flow cytometry. Graphs depict MFI data pooled from 6 independent experiments +SEM. Pre-specified groups were compared by a parametric unpaired T-test. **C.** Viable frequencies of MD4 lymph node B cells in culture at 72-hours post-stimulation with sHELD/T and pHELD/T in the presence of 20ng/ml BAFF assessed via live/dead stain applied to parent B220+ gate. Each datapoint is an independent sample from >= 4 independent experiments; lines connect samples from the same experiment. Groups were compared by a parametric paired T-test. **D.** As in 4A, but cells were cultured with IRAK1/4 inhibitor (1μM) and indicated stimuli (CpG 2.5 μM, pHELT-HD 1pM) for 3hrs and then stained to detect intracellular MYC expression. Histograms are representative of 2 independent experiments as well as an additional 2 experiments assessing MYC at 24hrs +/– MYD88. **E.** As in B, except either *Myd88*+/+ or *Myd88*–/– MD4 cells were cultured with either 2.5 μM CpG or 1pM pHELT-HD for 24hr. Histograms depict surface CD69 expression among live B220+ cells and are representative of 3 independent experiments. **F, G.** MD4 NUR77-eGFP reporter lymph node cells were labelled with CTV and cultured with indicated stimuli (1 μg/ml sHELD/T, 1pM pHELD/T) for 72 hours with 20ng/ml BAFF +/– 10ng/ml IL-4. Neighbouring histograms show division peaks, arrowheads denoting changes in division peak intensity between IL-4-treated and untreated samples. Representative plots depict CTV dilution (x-axis) relative to NUR77-GFP expression (y-axis) among live B220+ B cells assessed by flow cytometry. Shaded bars reference NUR77-GFP levels in dividing B cells stimulated with soluble Ag (sHELD/sHELT). (Right) Quantification of NUR77-GFP levels in the first division peak (cells that have undergone a single round of division). Data without IL-4 are representative of 5-7 independent experiments. Data comparing +/– IL-4 are representative of at least 4 independent experiments. **H.** As in F, but stained to detect intra-cellular C-MYC. Plots are representative of at least 3 independent experiments. **I. J.** Quantification of NUR77-GFP levels (I) and C-MYC (J) in the first division peak of polyclonal NUR77-eGFP reporter B cells cultured with anti-IgM 6.4μg/ml +/– 10ng/ml IL-4 as in S4E, H. Graph shows three biological replicates compared by parametric t-test. Data depicted are representative of > 6 independent experiments.

### pHEL reduces the NUR77-eGFP signaling threshold required for B cell proliferation in a manner that mimics T cell help

We previously showed that the NUR77-eGFP reporter reveals a sharp T cell receptor signaling threshold required for cell division^62^. Here we show that the reporter reflects an analogous threshold in B cells stimulated through the BCR and this is not altered by dose of anti-IgM, sHEL, or titration of the BTK inhibitor ibrutinib **(Supplementary Figure 4D)**. CD8 but not CD4 T cells exhibit a reduction in this threshold in response to IL-2 supplementation, entering cell cycle at lower levels of TCR-induced NUR77-EGFP expression^63^. Similarly, among dividing B cells supplemented with IL-4, the NUR77-eGFP level is markedly reduced **(Supplementary Figure 4E)**. Consistent with the hypothesis that pHEL may mimic co-stimulation, we find that the NUR77 level among dividing B cells stimulated with pHEL (even low density and low affinity) is markedly reduced about 10-fold compared to sHEL **(Figure 8F, G, Supplementary Figure 4F)**. Because this cannot be reproduced with any dose/affinity of sHEL or anti-IgM stimulation (**Supplementary Figure 4D, F**), this suggests that pHEL is delivering not only a more potent signal to B cells, but a qualitatively distinct one. The addition of IL-4 co-stimulation increases the number of divisions and reduces NUR77 levels among dividing cells responding to sHEL but does not alter the already low GFP expression following pHEL stimulation **(Figures 8F, G)**. This suggests that the pHEL BCR signal alone is sufficient to reduce the NUR77 signaling threshold for B cell proliferation and can mimic this effect of IL-4.

We previously reported that C-MYC marks an invariant threshold for proliferation in T cells that is enforced irrespective of co-stimulation^63^. Consistent with literature suggesting that MYC also imposes a minimal threshold for B cell proliferation^60, 64^, we find that both polyclonal and MD4 B cells entering cell division also express an invariant level of MYC, and neither pHEL nor co-stimulation alter this threshold in B cells **(Figure 8H-J, Supplementary Figure 4G, H)**. Rather, we postulate that both pHEL and co-stimulatory stimuli – relative to soluble Ag alone – induce and maintain MYC expression to a greater extent relative to NUR77– eGFP, and this is evident as early as 24 hrs before cell division has occurred (**Supplementary Figure 4B**). Efficient and sustained induction of MYC relative to NUR77 by pHEL – even in the absence of co-stimulation – may contribute to the T cell-independence of this mode of Ag display. By contrast, soluble Ag stimuli require co-stimulatory second signals to sustain MYC protein expression.

## Discussion

Although the assembled BCR complex has recently been visualized via cryo-EM, the mechanism by which Ag triggers signaling continues to be debated^12^. Yet, it remains widely accepted that multivalent Ags induce potent B cell responses by virtue of a high avidity interaction and receptor cross-linking. We reveal that this does not fully explain remarkable sensitivity of B cells to virus-like Ag display. Rather, we show that SVLS triggers signal amplification downstream of the BCR that is due, at least in part, to complete evasion of inhibitory co-receptor engagement but is independent of CD19 (see model, **Supplementary Figure 5**). This contrasts with membrane Ags that rely on CD19 for potency, and also with soluble Ags that engage ITIM pathways to curtail signaling and impose tolerance and T cell-dependence.

Evasion of inhibitory signaling by particulate Ag lowers the threshold for activation relative to soluble Ag, enabling B cells to sense and respond to *μ*M affinity epitopes. Reducing SVLS dose decreases the fraction of B cells that signal, but the amplitude of response is unaltered. This observation suggests that even a small number of particles with virus-like Ag display may be sufficient to trigger maximal signaling in a single B cell, and implies signal amplification downstream of BCR engagement by particulate Ag. It will be of great interest to rigorously identify the minimal number of SVLS and BCRs required to activate a B cell, and to determine whether signal amplification indeed occurs at the level PIP3 accumulation as biochemical evidence suggests.

Many biologically relevant Ags, including particulate Ag, are captured on APCs and presented directly to B cells *in vivo*^31–37^. Ag presented on a cell membrane dramatically reduces the threshold of B cell activation *in vitro*, enabling responses to very low-affinity Ags ^20–22^ and even to haptens that are non-stimulatory in soluble monovalent form^19^. Nevertheless, B cells can discriminate ligand affinity in this context in part via mechanical pulling forces at the contact interface ^14, 18, 23, 24, 65^. By contrast, we find no evidence of affinity discrimination to SVLS *in vitro* at the level of any proximal signaling readout assessed (including NF-*κ*B, MAPK, and calcium pathways). Indeed, maximal “all-or-none” responses on a single cell level to virus-like Ag display would be expected to preclude affinity discrimination. Moreover, such discrimination very early in the B cell response to bona fide virus is not merely dispensable, but undesirable^66^. Rather, later capture of virus by FDC to guide affinity maturation in the germinal center is a setting where affinity discrimination of membrane Ag is imperative and occurs^67^.

Imaging of B cells encountering membrane Ag has previously revealed progressive growth of BCR microclusters at the contact interface which spread and then contract into an immune synapse (IS) in a manner dependent on actin depolymerization. Such membrane re-organization can juxtapose BCRs to co-receptors such as CD19, while inhibitory molecules are excluded from the IS^20, 23, 27, 28^. Indeed, agents that depolymerize actin (to recapitulate such cell surface reorganization) are sufficient to trigger BCR signaling in a manner that is completely dependent upon CD19, IgM BCR, downstream signaling machinery such as PLC*γ*2, and is also constrained by inhibitory coreceptor CD22^28–30^. CD19 is the signaling component of the CD21/CD81/CD19 complex required to reduce the threshold for B cell activation by complement-coated Ag^55^. Yet, CD19 also has a unique, CD21-and complement-independent role in B cell activation; although it is dispensable for BCR signaling in response to soluble Ag, CD19 is required for sensitizing B cells to cell membrane Ag display^27, 29^. Distinct from membrane Ag, CD19 is dispensable for B cell response to particulate Ag suggesting this may not rely on actin dynamics to juxtapose coreceptors and BCRs. However, after particulate Ag is captured and presented on APCs *in vivo*^31–36^, B cell response is likely to become highly CD19-dependent both because of membrane-Ag presentation and complement fixation^55^.

Soluble Ag stimuli are highly constrained by LYN-dependent inhibitory pathways, but we show that SVLS entirely evades LYN-dependent inhibitory tone, resulting not only in heightened amplitude, but also markedly prolonged kinetics of signaling. Exclusion of CD22, Fc*γ*RIIb, and SHP1 has been visualized at the IS of B cells interacting with cell membrane-associated Ag, and enforced Fc*γ*RIIb engagement is sufficient to prevent BCR clustering in response to membrane Ag^20, 25, 26^. Yet, it remains to be determined whether these pathways are normally engaged at early time points by membrane Ag. Indeed, CD22 deletion comparably restricts calcium entry triggered by both soluble anti-kappa Ab stimulation and LatA-induced actin depolymerization^30^. LYN-deficient DT40 chicken B cell lines are completely defective in both membrane Ag and LatA responses^28, 68^, but unlike primary mouse B cells, DT40 express predominantly LYN SFK which is required for all ITAM signaling and does not isolate a role for ITIMs. It will be of great interest to define how LYN-dependent ITIM pathways impact membrane Ag signaling in mammalian B cells.

Importantly, we show that super-imposed stimulation with soluble antigenic stimuli (anti-IgM or even sHEL) is sufficient to prevent and/or terminate SVLS-induced signaling and this is at least partly LYN-dependent. This shows that forced engagement of LYN-dependent inhibitory signaling pathways can dominantly suppress pHEL responses. One candidate substrate of LYN that may mediate this effect is CD22 which can interact with the IgM BCR in a manner that depends on cis-ligands, and mediates inhibition by recruiting SHP1^48, 50, 57^. Indeed, engineered expression of CD22 ligands on HEL-conjugated lipid nanoparticles can fully abrogate BCR signaling, suppressing phosphorylation of AKT, CD19, as well as Erk, calcium entry, and I*κ*B degradation^69^. Whether CD22 alone or other ITIM-containing inhibitory co-receptors also account for LYN-dependent suppression of soluble but not SVLS signaling remains to be determined along with the effector PTPases required to do so (SHP-1, SHIP1 or others).

How do SVLS evade LYN-dependent inhibitory co-receptor engagement? Mature naïve B cells co-express IgM and IgD BCRs which not only exhibit markedly different abundance on the surface of naïve follicular B cells ^30^, but may also form separate pre-existing nanoclusters on resting cells prior to Ag encounter^29, 70^. Estimates for the size of these nanoclusters (60-240nm)^29, 70^ overlaps with the diameter of viruses found in nature. Such BCR nanoclusters harbor 30-120 molecules of IgD per cluster (corresponding to 6000-24000/µm^2^) and less densely packed IgM at the lower end of this range^29^. We postulate that aligning particle size and spacing of epitopes with that of BCR nanoclusters may maximize BCR engagement while also physically evading inhibitory co-receptor engagement. Indeed, pHELD-HD has 6,000 epitopes per µm^2^ which in turn corresponds roughly to 10nm optimal hapten spacing described in the “immunon” model and may trigger maximally potent B cell responses by even a small number of particles^8^. Naturally occurring viruses have epitope densities that range from 200 to 30,000 molecules per µm^2^; HIV occupies the lowest end of this spectrum and this may facilitate its immune evasion^39, 71^. Capacity to trigger potent signaling across a range of ED may also relate to conformational flexibility of SVLS. Future work could explore whether responses to rigidly structured VLPs and bona fide viral particles share features of SVLS signaling, and whether lipid nanoparticles offer unique advantages as scaffolds for flexible Ag display in vaccine design.

IgM and IgD BCRs also differ in sensitivity to different forms of Ag by virtue of distinct flexibility in their hinge regions; Ubelhart et.al suggested that monovalent Ag can signal through IgM but not IgD BCRs, while multivalent Ags can trigger both to signal^72^, although Sabouri et al. showed monomeric HEL can engage both^73^. High IgD expression on mature follicular B cells and preserved IgD on anergic B cells that downregulate IgM may sensitize these populations to particulate Ag display by viruses or virus-like particles^41, 42^. Differential IgM and IgD engagement by soluble and particulate Ag could also contribute to qualitative differences in signaling because IgM and IgD BCRs exhibit different co-receptor association – including CD22 and CD19^17, 29, 30^. These hypotheses warrant future investigation as does the responsiveness of other BCR isotypes expressed on memory B cells to SVLS.

Multivalent Ags – and viral Ag display in particular – can trigger T-independent B cell responses^1^. We show that multivalent Ag display on SVLS – independent of MYD88 or T cell co-stimulation – triggers not only strong MAPK and calcium pathway activation, but also robust NF-*κ*B activation (comparable to that induced by CD40 engagement), sustained MYC expression, as well as enhanced cell growth, survival and proliferation. Highly robust and bimodal NF-*κ*B activation by SVLS may reflect signal amplification downstream of PIP3 at the level of BTK dimerization and trans-autophosphorylation (Y322)^53^, as well as positive feedback over CARMA1/IKK*β*^74^. In addition, strong and prolonged calcium signals can activate IKK^46, 75^. Soluble Ag, by contrast, drives much less robust NF-*κ*B activation, triggers apoptosis, and exhibits a much higher NUR77-eGFP threshold for MYC-dependent proliferation, suggesting not only a quantitative but also a qualitative difference in the B cell response to soluble and particulate Ag. MYC is postulated to set an invariant threshold for proliferation (as we observe here)^64^, to drive cell growth, and to ‘fuel’ clonal expansion in the GC by orchestrating a transcriptional program of metabolic remodeling^60, 76, 77^. Robust and sustained NF-*κ*B and MYC expression may be sufficient to support pHEL-stimulated B cells to grow, survive, and proliferate in the absence of T cell input, and may help explain the molecular basis of T cell-independent B cell responses to multivalent Ags. Importantly, robust responses to SVLS are not restricted to MZ B cells which play critical roles in early response to T-independent multivalent model Ags ^44^. Rather, the biology we have uncovered is a general property of naïve, follicular B cells. Consistent with this, T-independent Ab responses are triggered by SVLS administered subcutaneously and without adjuvant *in vivo* in a manner likely to engage lymph node resident follicular B cells ^78^.

Autoreactive BCRs are common in the mature B cell repertoire, and one teleological explanation for their retention is to preserve a reservoir of protective specificities that can respond to cross-reactive viral epitopes under pressure to evolve towards self (near-self hypothesis^79, 80^). However, such B cells are constrained by IgM downregulation, dampened signal transduction (anergy), reduced survival, and suppression of CD86 upregulation – mechanisms that prevent the induction of autoimmunity *in vivo*^42, 45, 47^. Yet self-Ag arrayed on viruses and virus-like particles, including VSV, papillomavirus-like particles, Q*β* bacteriophage particles, and lipid nanoparticles can indeed break tolerance^9, 78, 81–84^. Previous work from us shows that high ED on lipid nanoparticles similar to those used in the present study is sufficient to drive secretion of class-switched IgG *in vivo* in response to self-Ag in the absence of T cell help^78^. How does epitope display on virus-like particles activate anergic B cells even in the absence of co-stimulation? The BCR signaling circuitry that enforces B cell anergy and dependence on “signal 2” (T cell-input) relies on ITIM-dependent inhibitory pathways^50^. We show that pHEL is sufficient to induce maximal calcium entry in a subset of MD4.ML5 anergic B cells and propose that SVLS break tolerance by selective evasion of LYN-dependent inhibitory tone. Differential engagement of such inhibitory tone may represent a fundamental mechanism by which self-reactive naïve B cells can accurately discriminate the same Ag presented in different forms, enabling anergic cells to remain unresponsive in the face of soluble self-Ag until recruited into an immune response by bona fide viral encounter.

Together, these molecular insights reveal that density of epitope display on particles of viral size – independent of other viral features like encapsulated nucleic acid – serves as a stand-alone danger signal sufficient to breach B cell anergy by triggering a unique mode of BCR signaling that differs from both typical soluble and membrane Ags. We identify a role for inhibitory coreceptors that confers potency to such particles independent of avidity. Addressed in independent work (biorxiv preprint 2023)^59^ is understanding how nucleic acid delivered by SVLS or viruses activate B cells synergistically^85–87^. Our data suggests that mature B cells are intrinsically ‘built’ as a sensor for virus-like supramolecular structures and poised to mount rapid, potent, and initially T-independent responses to viral threats.

## Materials and Methods

### Mice

NUR77-EGFP and IgHEL (MD4) mice were previously described^41, 42^. C57BL/6 and CD45.1+ BoyJ mice were originally from The Jackson Laboratory. Lyn^−/–^ mice were previously described^56^. MYD88^tm^^1.Defr^ (Jackson labs #009088) (MYD88^−/–^ herein) encodes a deletion of exon 3 of the myeloid differentiation primary response gene 88 gene^88^. Cd19^tm^^1(Cre)Cgn^ (Jackson labs #006785) (CD19^−/–^ herein) has a Cre recombinase gene inserted into the first exon of the CD19 gene and thus abolishing Cd19 gene function^89^. ML5 mice (Jackson Labs #002599) that expresses sHEL Tg were previously described^42^. All strains were fully backcrossed to the C57BL/6J genetic background for at least 6 generations. Mice were used at 6-9 wk of age for all functional and biochemical experiments. All mice were housed in a specific pathogen-free facility at University of California, San Francisco, according to the University Animal Care Committee and National Institutes of Health (NIH) guidelines.

### SVLS synthesis and quantification

All SVLS used throughout this study were prepared following protocols previously described by us^38^. In brief, SVLS were constructed using nonimmunogenic lipids, with phosphatidylcholine and cholesterol comprising ≥ 95% of all lipids. We prepared unilamellar liposomes using a mixture of DSPC, DSPE-PEG maleimide, and cholesterol. Empty liposomes were generated using PBS and extruded membrane with a pore size of 100 nm. Hen egg lysozyme (HEL) recombinant proteins with free engineered cysteines were conjugated to the surface of liposomes via maleimide-thiol chemistry at specific engineered cysteine and programmable epitope density by modulating molar percentage admixed DSPE-PEG maleimide (0.5-5%). A large excess of free cysteines at the end of a 1 h cross-linking reaction to quench all of the available maleimide groups that remain on the liposomal surface. After this conjugation, the liposomes were purified away from free excess HEL proteins by running through a size exclusion column (SEC). Epitope density was quantified using methods that we established previously^38, 90^ that were also validated by single-molecule fluorescence^91, 92^. Control SVLS (pLIP02) had no surface protein conjugation. Recombinant HEL-WT, HEL-D (R73E, D101R), HEL-T (R73E, D101R, R21Q) proteins were overexpressed in *E. coli* and purified to >95% purity following our established protocols^38^ and are used throughout except WT HEL from Sigma (1 *μ*g/ml) is used in S2F, 4C, 4D, 7I, and 8B.

### Antibodies and Reagents

#### Antibodies for surface markers

Streptavidin (SA) and antibodies to B220 (RA3-6B2), CD19 (1D3), CD21 (7G6), CD23 (B3B4), CD86 (GL-1), CD69 (H1.2F3), CD45.1 (A20), CD45.2 (104) conjugated to biotin or fluorophores (BioLegend, eBiosciences, BD, or Tonbo).

#### Antibodies for intra–cellular staining

pErk1/2 T202/Y204 mAb (clone 194G2, cat. 4377S) pS6 S235/236 mAb (clone 2F9, cat. 4856S), I*κ*B*α* Ab (cat. 9242S), NF-κB p65 mAb (clone D14E12, cat. 8242S), C-MYC mAb (clone D84C12, cat. 5605S) were from Cell Signaling Technologies; Donkey anti–rabbit secondary Ab conjugated to APC was from Jackson Immunoresearch.

#### Antibodies for immunoblots

All antibodies except where otherwise noted were from Cell Signaling: C-MYC mAb (clone D84C12), pERK T202/Y204 mAb (clone 197G2, cat. 4377S), GAPDH Ab (14C10, cat. 3683s), pCD19 Y531 Ab (cat. 3571S), pSHP-1 Y564 mAb (clone D11G5, cat. 8849S), pSHIP1 Y1020 Ab (cat. 3941S), pPLC*γ*2 Y1217 Ab (cat. 3871S), pCD79A Y182 Ab (cat. 5173S), pSYK Y525/526 mAb (clone C87C1, cat. 2710S); pBTK Y223 mAb (clone D9T6H, cat. 87141S), pPI3K [p85 Y458, p55 Y199] mAb (clone E3U1H, cat. 17366S), pAKT T308 mAb (clone D25E6, cat. 13038S), pAKT S473 Ab (cat. 9271S), pIKK*α/b* S176/180 mAb (clone 16A6, cat. 2697S) were from Cell Signaling Technology, anti-phosphotyrosine clone 4G10 was made from a hybridoma and was a gift from the Weiss lab. pCD22 Y822 [mouse Y837] mAb (clone Y506, cat. ab32123) was from Abcam. Anti-mouse-and anti-rabbit-HRP secondary Ab were from Southern Biotech.

#### Stimulatory Antibodies and reagents

Goat anti-mouse IgM F(ab’)2 was from Jackson Immunoresearch; Murine IL-4 (10ng/ml, cat#214-14, Peprotech), anti-CD40 (1*μ*g/ml, hm40-3 clone; BD Pharmingen), recombinant murine BAFF (20ng/ml, cat#2106-BF, R&D), LPS (10*μ*g/ml O26:B6; Sigma), CpG (2.5*μ*M ODN 1826; InvivoGen).

#### Media

Complete culture media was prepared with RPMI-1640 + L-glutamine (Corning-Gibco), Penicillin Streptomycin L-glutamine (Life Technologies), 10 mM HEPES buffer (Life Technologies), 55 mM *β*-mercaptoethanol (Gibco), I mM sodium pyruvate [(Life Technologies), Non-essential Amino acids (Life Technologies), 10% heat inactivated FBS (Omega Scientific). This was used for all *in vitro* culture and stimulation except for calcium signaling experiments. Calcium flux media was prepared with RPMI, HEPES, PSG as above and 5% FCS.

#### Inhibitors

SYKi (Bay 61-3606 1*μ*M) and PI3Ki (Ly290049 10*μ*M) were from Calbiochem, BTKi (ibrutinib 100nM from Jack Taunton, UCSF), IRAK1/4i (R568 1*μ*M, gift from Rigel)

### Flow Cytometry and data analysis

After staining, cells were analyzed on a Fortessa (Becton Dickinson). Data analysis was performed using FlowJo (v9.9.6 and v10) software (Treestar Inc.). Proliferative indices ‘division index’ and ‘% divided’ were calculated using FlowJo. Statistical analysis and graphs were generated using Prism v6 (GraphPad Software, Inc). Statistical tests used throughout are listed at the end of each figure legend. Student’s paired or unpaired T test (two-tailed and parametric) was used to calculate p values for all comparisons of two pre-specified groups depending on experimental design. ANOVA was performed when more than two groups were compared to one another. Error bars in graphs represent SEM.

### Intracellular staining to detect pERK, pS6, IκB, and cMYC

Following *in vitro* stimulation for varying times, 10^6^ cells / well were resuspended in 96-well plates, stained with fixable live/dead dye as above for overnight time points, fixed in 2% paraformaldehyde (PFA) for 10 min, washed in FACS buffer, and permeabilized with ice-cold 90% methanol at −20 °C overnight, or for at least 30 minutes on ice. Cells were then washed in FACS buffer, stained with intra-cellular Ab x 45 minutes, washed in FACS buffer, and then stained with Alexa 647-conjugated secondary Goat anti-Rabbit antibody along with directly conjugated antibodies to detect surface lineage and/or subset markers for 45 minutes. Samples were washed and then refixed with 2%PFA.

### Intracellular Calcium Flux

Cells were loaded with 5 μg/mL Indo-1 AM per manufacturer’s instructions (Life Technologies) and stained with lineage markers B220, CD23, CD21 for 15 min. Cells were rested at 37 °C for 3 min, and Indo-1 fluorescence was measured by FACS immediately prior to and after stimulation to determine intracellular calcium.

### Live/dead staining

LIVE/DEAD Fixable Near-IR Dead Cell Stain kit (Invitrogen). Reagent was reconstituted as per manufacturer’s instructions, diluted 1:1000 in PBS, and cells were stained at a concentration of 2 × 10^6^ cells /100*μ*L on ice for 10 min.

### Vital dye loading

Cells were loaded with CellTrace Violet (CTV; Invitrogen) per the manufacturer’s instructions except at 5 × 10^6^ cells/ml rather than 1 × 10^6^ cells/ml.

### In Vitro B Cell Culture and Stimulation

Splenocytes or lymphocytes were harvested into single cell suspension through a 40*μ*m cell strainer, subjected to red cell lysis using ACK (ammonium chloride potassium) buffer in the case of splenocytes, +/– CTV loading as described above, and plated at a concentration of 2.5-10 × 10^5^ cells/200 *μ*L (depending on assay) in round bottom 96 well plates in complete RPMI media with stimuli for 1-3 days. *In vitro* cultured cells were stained to exclude dead cells as above in addition to surface or intra-cellular markers for analysis by flow cytometry depending on assay.

### Immunoblot analysis

Purified splenic/LN B cells (protocol described below) were harvested from mice, pre-warmed for 15 min at 37°C, and mixed with stimuli in RPMI for 3 min at 37°C. Following stimulation, cells were lysed with 1% NP40 lysis buffer with protease and phosphatase inhibitors for 15 min at 4°C, and centrifuged for 10 min at 20,000g to remove cellular debris. The supernatants were denatured at 95^°^C for 5 min in SDS sample buffer with 2.5% *β*-ME. Lysates were run on Tris-Bis thick gradient (4%-12%) gels (Invitrogen) in MOPS buffer, and transferred to PVDF membranes with a Mini-Protean Tetra cell (Bio-rad). Membranes were blocked for 1 h with 3% BSA TBST, and then probed with primary antibodies listed above, overnight at 4^°^C. The next day, membranes were washed three times with TBST for 10 min, and incubated with HRP-conjugated secondary antibodies. Blots were developed utilizing a chemiluminescent substrate (Western Lightning Plus ECL, Perkin Elmer), and visualized with a ChemiDoc Touch Imaging System (Bio Rad). Quantification of western blots was performed with Image Lab Software (Bio Rad).

### B cell purification

B cell purification was performed utilizing MACS separation per manufacturer’s instructions. In short, pooled spleen and/or lymph nodes were prepared utilizing the B cell isolation kit (Miltenyi), and purified by negative selection through an LS column (Miltenyi).

### Nuclei isolation / nuclear staining

The nuclei isolation protocol was based on previously published report^93^.

### Nuclei isolation

In brief, purified splenic/LN B cells (protocol described below) were harvested from mice and pre-warmed in a 96-well plate for 10 min at 37°C with 1.5x10^6^cells/100 μl per well in complete media (RPMI with 10% FBS, penicillin, streptomycin, glutamine, HEPES, *β*-ME, sodium pyruvate, and non-essential amino acids). Cells were stimulated with 100 μl of 2x stimuli for 20 min at 37°C. Stimulated cells were spun at 300 x *g* and 4°C, and the pellets were immediately resuspended with 250 μl of ice-cold Buffer A containing 320 mM sucrose, 10 mM HEPES (Life Technologies), 8 mM MgCl2, 1× Roche EDTA-free complete Protease Inhibitor, and 0.1% (v/v) Triton X-100 (Sigma-Aldrich). After 15 min on ice, the plate was spun at 2000 × *g* and 4°C for 5 min. This was followed by two 250 μl washes with Buffer B (Buffer A without Triton X-100) and spinning at 2000 × *g* and 4°C for 5 min. After the final wash, pellets were resuspended with 200 μl Buffer B containing 4% paraformaldehyde (electron microscopy grade; Electron Microscopy Sciences), and nuclei were rested on ice for 30 min for fixation. The nuclei were spun at 2000 x *g* and 4°C for 5 min, and followed by two washes and resuspension in FACS Buffer (1× PBS with 2% FBS and 8mM MgCl2), centrifuging at 1000 × *g* and 4°C for 5 min to sufficiently pellet nuclei.

### Nuclei staining

Isolated nuclei were washed with 200 μl Perm Buffer (FACS Buffer with 0.3% Triton-X 100) and spun at 1000 × *g* and 4°C for 5 min. Isolated nuclei were stained with 50 μl purified rabbit anti-mouse NF-κB p65 (Cell Signaling 8242S) diluted (1:100) in Perm Buffer for 30 min on ice. The 96-well plate was spun at 1000 x *g* and 4°C for 5 min, and washed with 200 μl FACS Buffer. The nuclei were then stained with 50 μl anti-rabbit IgG-APC (Jackson Immunoresearch, 711-136-152) diluted (1:100) in Perm Buffer for 30 min on ice in the dark. The nuclei were spun at 1000 x *g* and 4°C for 5 min, and washed with 200 μl FACS Buffer. Nuclei were resuspended in 200 μl FACS Buffer and analyzed immediately on a BD Fortessa cytometer.

## Acknowledgments

We thank our funders – R01AI155653 (WC, JZ), NIAID R01AI148487 (JZ), NIAMS R01AR069520 (JZ), NIAID AAI Postdoctoral Fellowship (JFB and JZ), NIH Training Grant T32 AI007334 (JR). We thank Dr. Arthur Weiss for helpful discussions and critical reading of the manuscript. We thank Al Roque for help with mouse husbandry.

---

**Supplementary Figure 1.**
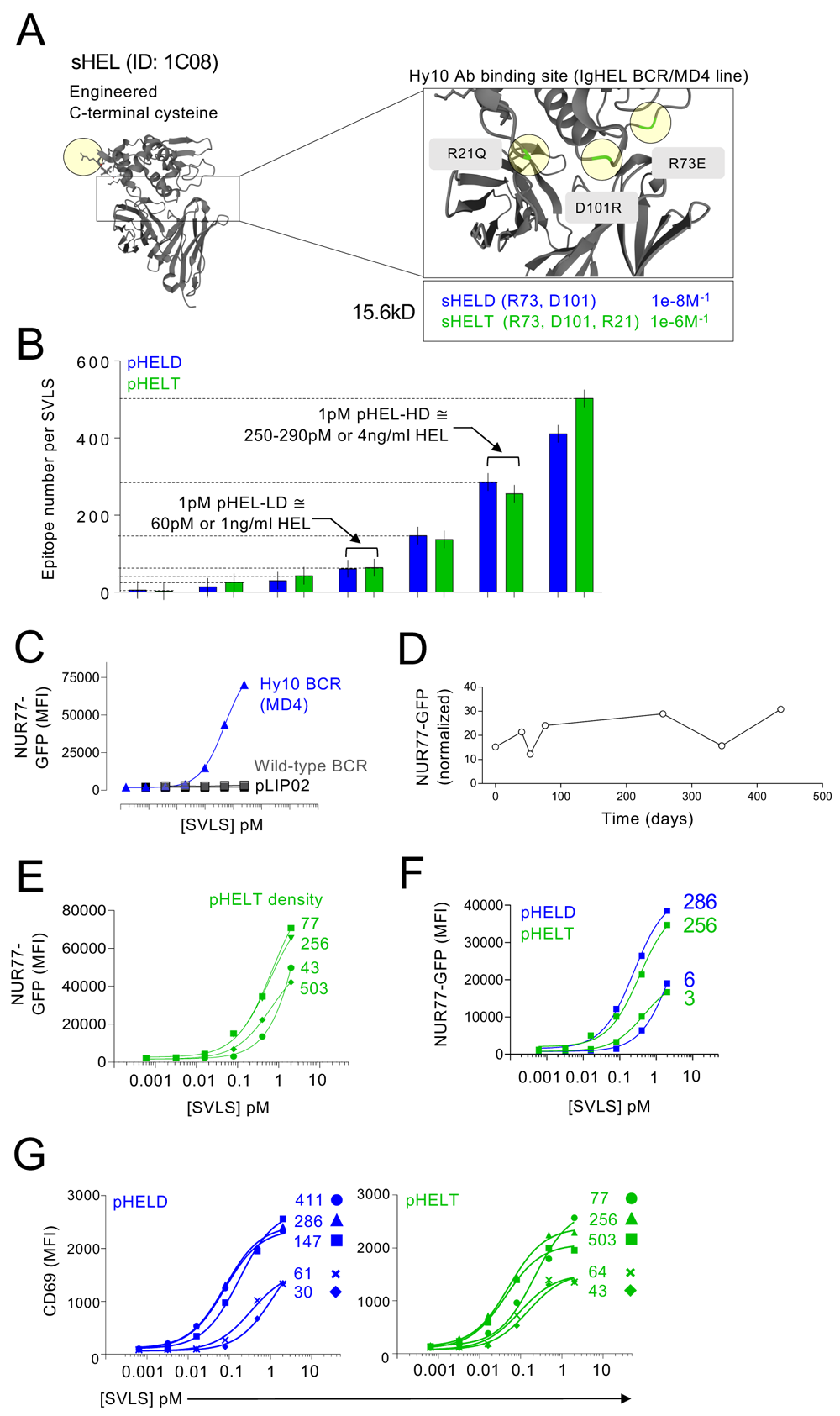
Associated with Main **Figure 1**. **A.** Hen egg lysozyme (HEL) was engineered with a c-terminal cysteine residue (yellow shadowing) that preserves native protein structure but allows stable conjugation in a fixed orientation to SVLS via maleimide chemistry. Point mutations (zoomed box) were also introduced into sHEL epitope which is bound by the Hy10 BCR and were previously described to reduce binding affinity^38^. HELD carrying two mutations or HELT carrying three mutations, along with the cysteine modification, were then purified from an overexpression system in E. coli. **B.** A library of SVLS was constructed with purified HELD or HELT conjugated at various densities to liposomes yielding pHELD/T particles with 3-500 antigenic epitopes per particle. Epitope density is calculated as [HEL]/[lip] in M. For pHELD/T with high ED 250-290, 1pM pHEL = 250-290nM HEL (∼4ng/ml; 15.6kD HEL mol weight). **C.** Congenically-marked MD4 (Hy10) and wild-type splenocytes were stimulated with pHELD or a naked control particle (pLIP02). NUR77-GFP reporter expression along with congenic markers CD45.1/2 was analysed in B220+ cells 24 hours later as in Main Fig 1C. Axes were intentionally off-set to visualize lack of response to pLIP02/in wild-type B cells. **D.** NUR77-GFP reporter expression was analysed in MD4 B cells at 24 hours post-stimulation with the same batch of pHELD at various dates after its synthesis. Reporter expression was normalized to the pLIP02 condition in each experiment. **E.** As in main Fig 1E but with pHELT instead of pHELD at varying epitope densities. Data is from a single experiment and representative of at least 3 independent experiments. **F.** As in E, but stimulated with SVLS carrying high or ultra-low density of HELD or HELT. Data is from a single experiment. **G.** As in E, but CD69 was analysed at 24 hours post-stimulation with pHELD (left) or pHELT (right). Data are from a single experiment and representative of at least 3 independent experiments. Curves were modelled with three-parameter nonlinear regression (C, E, F, G).

**Supplementary Figure 2.**
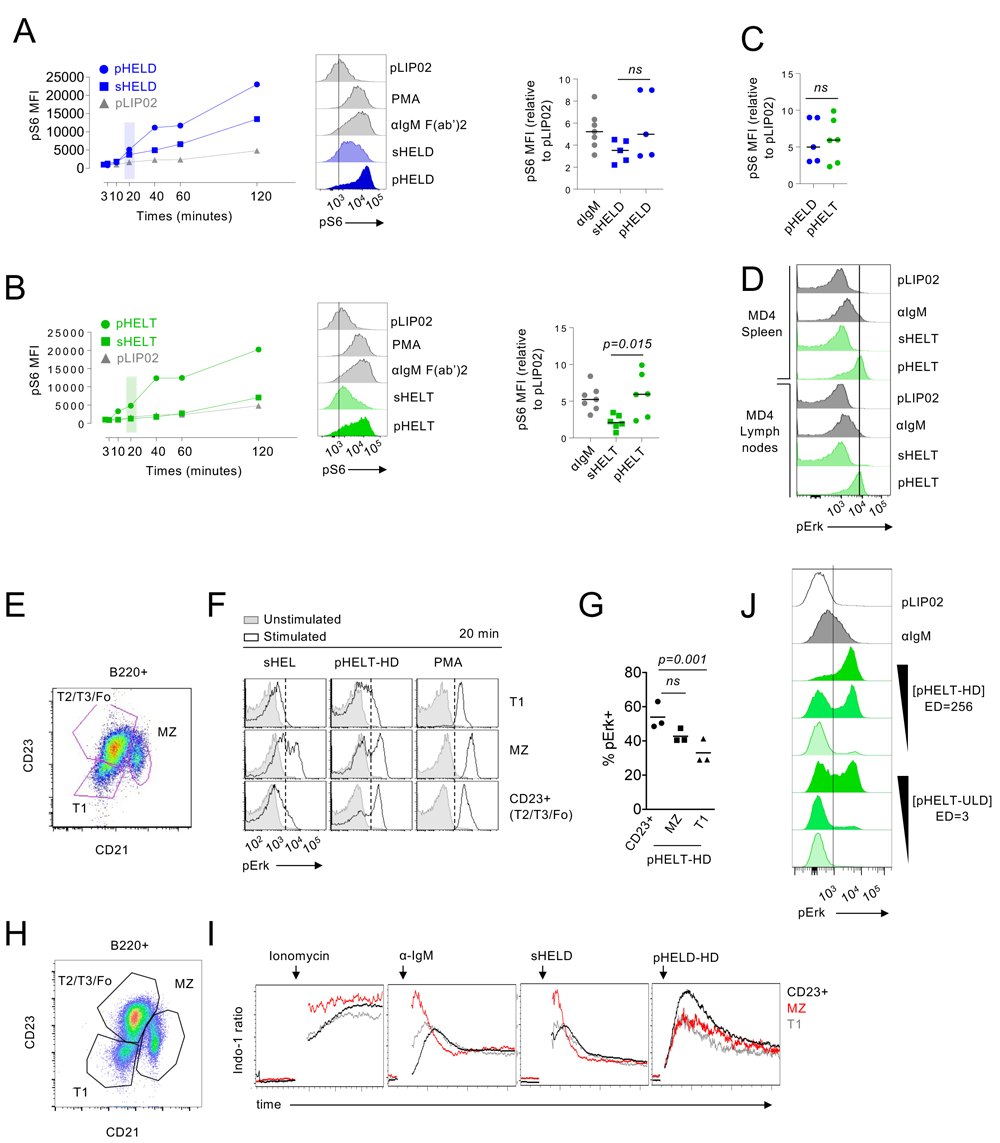
Associated with Main **Figure 2**. **A, B.** Pooled splenocytes and lymphocytes from MD4 mice were stimulated and stained as in Main Figure 2A, B but assessed for intra-cellular pS6 over a time course. Graphs (left) and histograms (middle) depict pS6 in B220+ cells from a single time course and is representative of at least 2 independent experiments. Graphs (right) depict B cell pS6 MFI after 20 min stimulation; each point represents an independent experiment and data are pooled >= 5 experiments depending on stimulus. **C.** Graph depicts pHELT and pHELD data from A, B graphed together for comparison. Pre-specified groups in A-C were compared by a paired parametric T-test. **D.** As in 2A, B but comparing pErk in stimulated B220+ splenocytes and lymph node cells separately at 20 mins. Histograms are from at least 2 independent experiments. **E-G**. As in 2A, but splenocytes were stained with B220, CD21, and CD23 and gated to identify pErk expression in B220+ subsets (E): immature T1 (CD21negCD23neg), marginal zone (CD21hiCD23int) and T2/T3/Follicular mature (CD23+CD21int). Representative histograms show pErk in gated subsets following sHEL-WT 1 μg/ml, 1pM pHELT-HD (ED 256), or PMA stimulation (F). Quantification of %pErk+ cells from three independent experiments (G). Groups were compared by one-way ANOVA. Data in E-G are representative of at least 8 independent experiments. **H, I.** Pooled MD4 splenocytes and lymphocytes were loaded with Indo-1 calcium indicator dye, stained with surface markers to detect B220+ subsets (H) as in S2E, and stimulated with anti-IgM (10 μg/ml), sHELD (1 μg/ml), or pHELD-HD (1pM). Calcium entry was assessed by flow cytometry for at least 3 minutes. Ratio of bound/unbound Indo-1 fluorescence over time is plotted (I). Data are representative of at least 4 independent experiments. **J.** As in Main Figure 3A, B except CD23+ B cells stimulated with pHELT-HD ED 256 and pHELT ultra low ED 3 are compared at 10, 1, 0.1pM doses. Data are representative of 4 independent experiments.

**Supplementary Figure 3.**
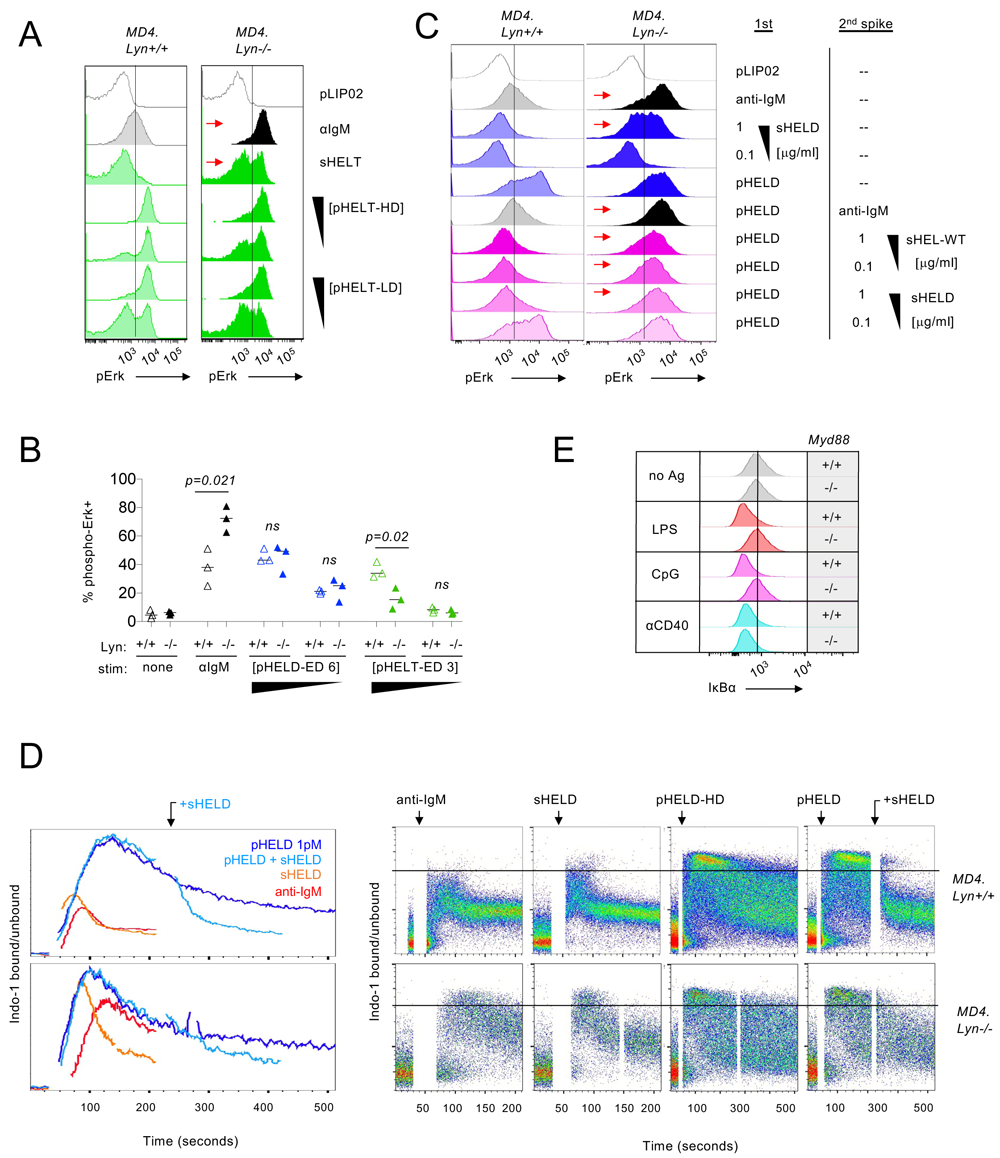
Associated with Main **Figure 6**. **A.** As in Main Figure 6B, but with sHELT and pHELT (HD/LD 256, 64) stimuli. Data are representative of 4 independent experiments. **B.** As in Main Figure 6B, C, except pHELD ultra low ED 6 and pHELT ultra low ED 3 are used at 10pM and 1pM doses. Graph depicts data from 3 independent experiments. Pre-specified groups were compared by parametric T-tests and are plotted together on the same graph for convenience. **C.** As in Main Figure 6E, but with inclusion of anti-IgM 10 μg/ml as second stimulus following pHELD. Data are representative of 4 independent experiments. **D.** As in Main Figures 6F, G but with sHEL 1 μg/ml added in at peak calcium response following pHEL stimulation. Data are representative of at least 3 independent experiments. **E.** As in Main Figure 7I, but with LPS 10μg/ml, CpG 2.5μM, or anti-CD40 1μg/ml stimulation. Data are representative of 3 independent experiments.

**Supplementary Figure 4.**
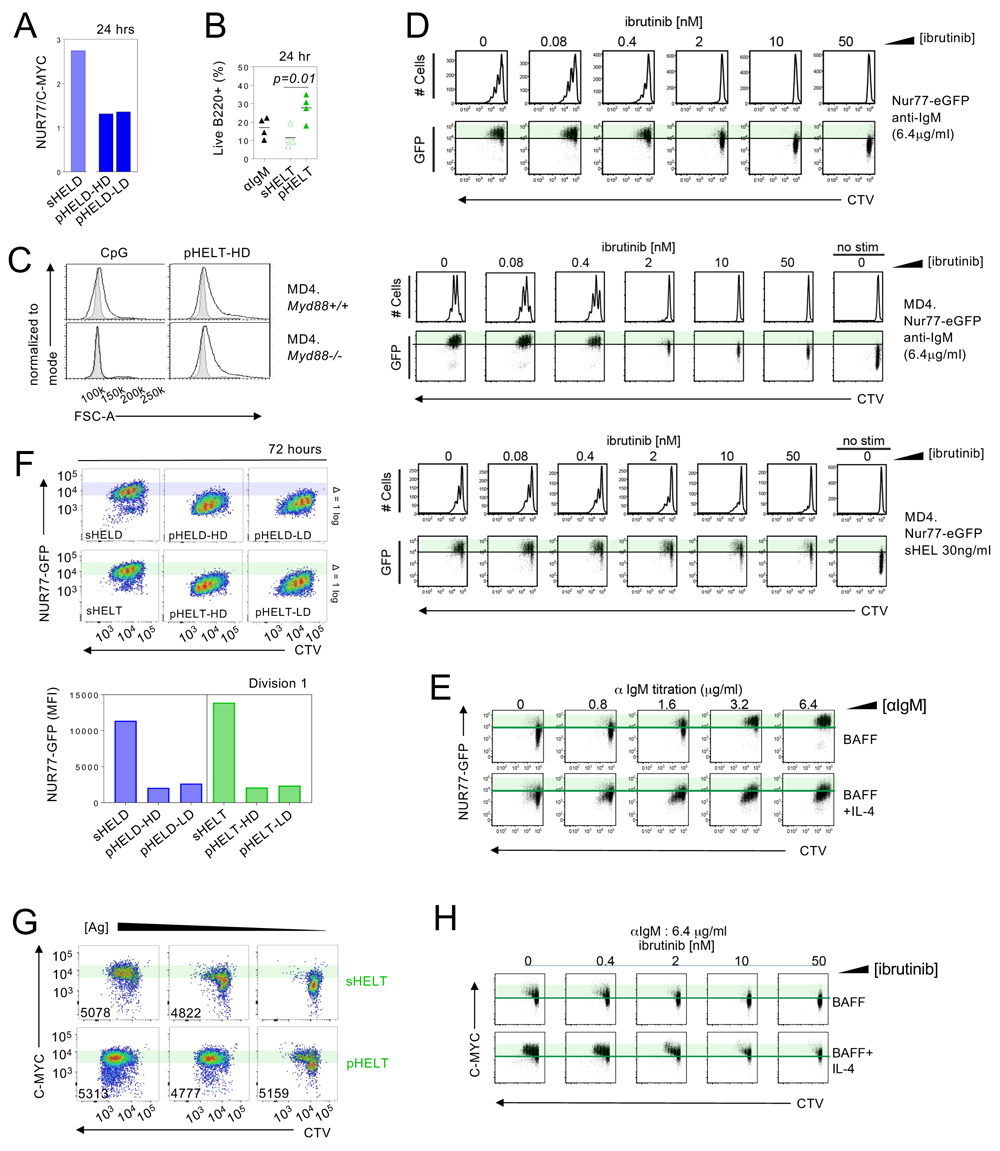
Associated with Main **Figure 8**. **A.** Pooled splenocytes and lymphocytes from MD4 NUR77-eGFP reporter mice were cultured for 24 hr with sHELD 1μg/ml or pHELD-HD, LD (1pM), fixed, permeabilized, stained and analysed for GFP and intra-cellular C-MYC. A ratio of NUR77 and C-MYC among B220+ cells was calculated. Data plotted is from a single experiment and is representative of at least 3 independent experiments. **B.** MD4 pooled splenocytes and lymphocytes were cultured for 24 hr with anti-IgM 10μg/ml, sHELT 1μg/ml or pHELT-HD 1pM (without supplemental BAFF). Viable frequencies of MD4 B cells were assessed via live/dead stain. Each datapoint is an independent biological replicate. Groups were compared by a parametric T-test. **C.** As in Main Fig 8E except histograms depict FSC-A among live B220+ cells as a measure of cell growth and are representative of 4 independent experiments. **D.** NUR77-eGFP reporter splenocytes (top) or MD4.NUR77-eGFP splenocytes (middle and bottom) were loaded with vital dye CTV and cultured for 72 hours in the presence of either 6.4μg /ml anti-IgM or 30ng/ml sHEL (bottom), supplemented with 20ng/ml BAFF, and various concentrations of ibrutinib (0-50nM), and stained to detect viability and B220. Plots depict CTV and GFP expression in live B220+ B cells. Data are representative of at least 3 independent experiments. **E.** NUR77-eGFP reporter splenocytes cultured in the presence of 20ng/ml BAFF as in D except with varying doses of anti-IgM +/-10ng/ml IL-4. Plots depict CTV and GFP expression in B220+ B cells. Data are representative of at least 6 independent experiments. **F.** As in Main Fig 8F, G MD4 NUR77-eGFP lymphocytes were cultures with stimuli + 20ng/ml BAFF for 72 hrs: 1 μg/ml sHELD/T, 1pM pHELD/T-HD or -LD. These data correspond to those depicted in Fig 8F, G but now include plots and quantification for pHELD/T-LD for direct comparison. Representative plots depict CTV dilution (x-axis) relative to NUR77-GFP expression (y-axis) among live B220+ B cells assessed by flow cytometry. Shaded bars reference NUR77-GFP levels in dividing B cells stimulated with soluble antigen (sHELD/sHELT). (Bottom) Quantification of NUR77-GFP levels in the first division peak (cells that have undergone a single round of division). Data are representative of 5-7 independent experiments. **G.** As in Main Fig 8H, but comparing intra-cellular C-MYC expression among live CTV-loaded B220+ B cells after 72hr culture across a range of doses: pHELT (1, 0.1, 0.01pM) and sHELT (500, 50, 5 ng/ml). Inset values represent MFI of division 1 cells in each plot. Data are representative of 3 independent experiments. **H.** As in E, but cultured with fixed 6.4 μg/ml dose of anti-IgM and titration of ibrutinib, and comparing C-MYC among live CTV-loaded B220+ B cells after 72hr culture. Data are representative of at least 6 independent experiments.

**Supplementary Figure 5.**
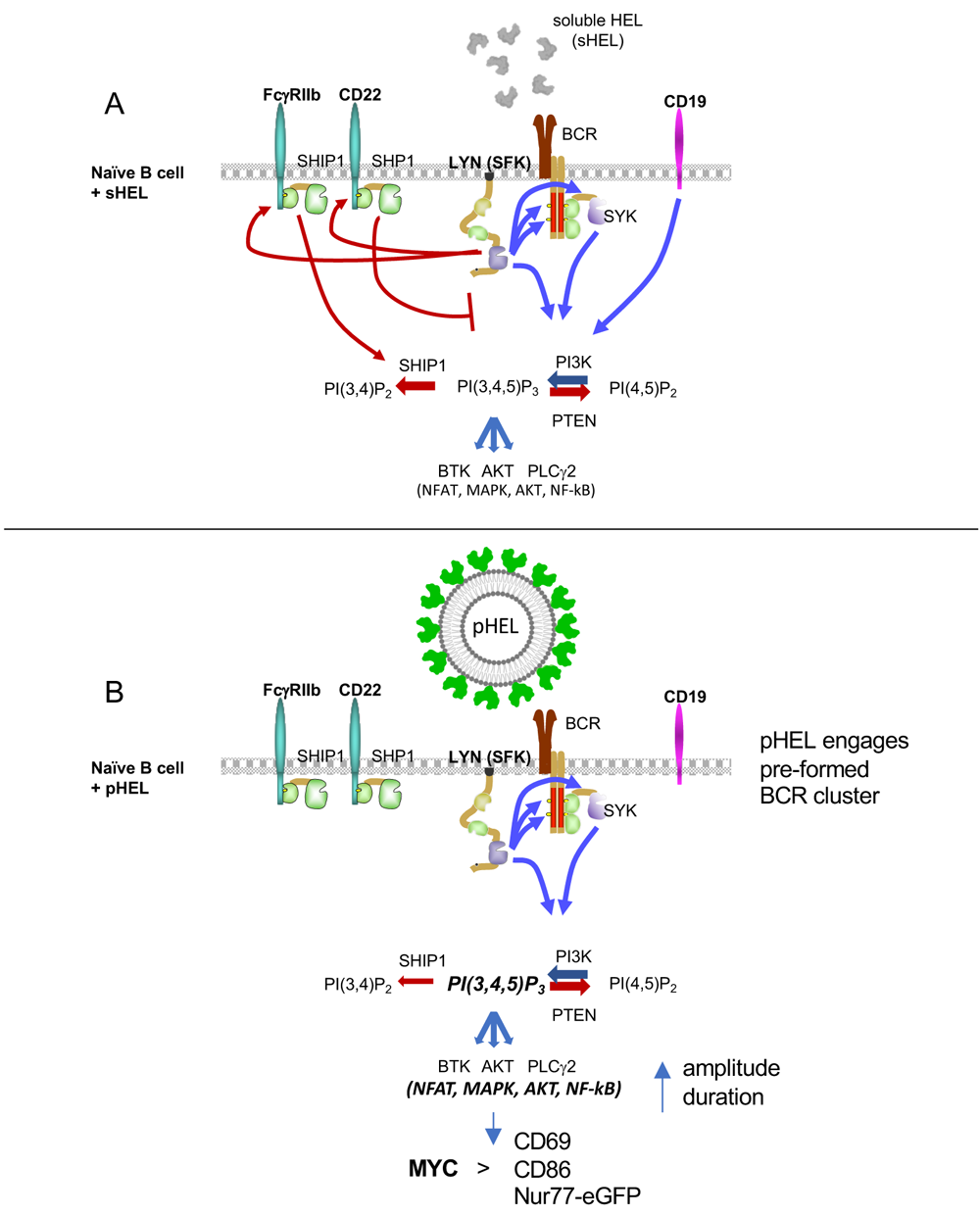
Model. **A.** Signalosome assembly in naïve B cells upon BCR stimulation by soluble Ag. BCR signal transduction requires sequential action of Src family kinases (SFKs) and SYK kinase. CD19 engagement amplifies PI3K activation. While multiple SFKs can mediate ITAM signaling downstream of the BCR, the SFK LYN plays a non-redundant role in phosphorylating ITIM-containing inhibitory coreceptors which in turn recruit PTPases SHP1 and SHIP1 that suppress PIP3. Dynamic regulation of PIP3 at the plasma membrane controls amplitude of signaling by recruiting downstream mediators including AKT, BTK, and PLCγ2 to orchestrate transcriptional programs mediated by NFAT, NF-κB and other factors. **B.** SVLS with appropriately spaced epitopes robustly engage pre-existing BCR nanoclusters, but evade co-inhibitory receptors resulting in downstream signal amplification. SVLS do not rely upon CD19 engagement for signal amplification in vitro. In the absence of inhibitory PTPase engagement via ITIM-containing inhibitory receptors, PIP3 accumulates at the plasma membrane leading to enhanced and prolonged signalosome assembly and activity. Robust NFAT and NF-κB accumulation in the nucleus and AKT-dependent signals mimic co-stimulation and promote MYC expression, resulting in T-independent cell growth, survival and proliferation.

